# betAS: intuitive analysis and visualisation of differential alternative splicing using beta distributions

**DOI:** 10.1101/2022.12.26.521935

**Authors:** Mariana Ascensão-Ferreira, Rita Martins-Silva, Nuno Saraiva-Agostinho, Nuno L. Barbosa-Morais

## Abstract

Next generation RNA sequencing allows alternative splicing (AS) quantification with unprecedented resolution, with the relative inclusion of an alternative sequence in transcripts being commonly quantified by the proportion of reads supporting it as percent spliced-in (PSI). However, PSI values do not incorporate information about precision, proportional to the respective AS events’ read coverage. Beta distributions are suitable to quantify inclusion levels of alternative sequences, using reads supporting their inclusion and exclusion as surrogates for the two distribution shape parameters. Each such beta distribution has the PSI as its mean value and is narrower when the read coverage is higher, facilitating the interpretability of its precision when plotted. We herein introduce a computational pipeline, based on beta distributions accurately modelling PSI values and their precision, to quantitatively and visually compare AS between groups of samples. Our methodology includes a differential splicing significance metric that compromises the magnitude of inter-group differences, the estimation uncertainty in individual samples, and the intra-group variability, being therefore suitable to multiple-group comparisons. To make our approach accessible and clear to both non-computational and computational biologists, we developed `betAS`, an interactive web app and user-friendly R package for visual and intuitive differential splicing analysis from read count data.

## INTRODUCTION

Species with similar numbers of protein-coding genes can have markedly discrepant numbers of functional proteins (1). Alternative splicing (AS) greatly amplifies transcriptomic diversity through the production of several RNA isoforms from a single gene. Virtually all human protein-coding genes undergo AS (2), that expands the potentially functional transcriptome (1) and contributes to the pre-translation regulation of gene expression (3).

Developmental stage- and tissue-specific transcriptomes are fine-tuned through the highly regulated usage of AS variants (4) and perturbations in the physiological regulation of tissue-specific AS have been reported in disorders with different aetiologies, such as autism (5), myotonic dystrophy (6) and cancer (7). Moreover, splicing modulation has been therapeutically used to direct AS towards protective isoforms. Therefore, soundly quantifying AS and differences in splicing between conditions not only helps to elucidate molecular bases of tissue physiology, but can also reveal therapeutic targets (8, 9).

High-throughput sequencing of RNA (RNA-seq) allows unprecedented precision in AS quantification (2). Computational tools developed for the identification and quantification of differentially spliced genes from RNA-seq data can be classified into isoform-resolution or count-based approaches (10, 11). While isoform-resolution methods, also called “multi-read” models (10), aim at directly estimating the relative abundances of full-length transcripts in each sample (11, 12), count-based methods aim at detecting differential usage of local features (e.g., exons) rather than the whole gene, based on the number of RNA-seq reads unambiguously assigned to each unit (10–12). Counting units refer to splicing events, i.e., gene regions showing local variation at the mRNA level due to two possible splicing outcomes, such as the inclusion or skipping of an alternative exon (13). A widely used metric for quantifying the fraction of mRNA isoforms that include an alternative sequence is the percent or proportion spliced-in (PSI), given by the ratio between inclusion and the sum of inclusion and skipping reads (14). For instance, AS quantification programmes `rMATS` (15), `vast-tools` (16), `Whippet` (17), `SUPPA2` (18), `MAJIQ` (19) and `MISO` (20) use PSI or analogous metrics for quantifying inclusion of alternative sequences.

Most differential AS analysis tools build on knowledge from differential gene expression analysis and apply linear modelling to define differentially spliced events between two or more groups (`limma` (21), `edgeR` (22), `DEXSeq` (23), `JunctionSeq` (24), `PennDiff` (25) and `DJExpress` (26)), while others quantify the difference of inclusion levels based on geometric distances between gene-level vectors (`SplicingCompass` (27), `DIEGO` (28)). However, most of these approaches have limited applicability to the typical small sample sizes used in biomedical research. `MISO`’s differential splicing approach relies on a probabilistic model that applies Bayesian statistics to isoform expression levels and thereby provide confidence intervals for PSI estimates for individual samples. However, `MISO` does not support multiple biological replicates directly.

Differential splicing analyses with small sample sizes should be supported by a compromise between modelling the uncertainty in the estimation of inclusion levels of AS events in individual samples and accounting for the biological variability among replicates. Although most tools overlook this compromise, `rMATS`’ (15) reportedly robust quantification pipeline (11) implements it by modelling: a) the uncertainty in the estimate of alternative sequence inclusion in individual replicates as a function of the total number of supporting RNA-seq reads, assuming these follow a binomial distribution; and b) the variability of such inclusion level across biological replicates as random effects in a mixed model. Similarly, `vast-tools` (16), `Whippet` (17), and `MAJIQ` (19) use the beta distribution to model the PSI variance in a read coverage-dependent way. `SUPPA2` (18) also incorporates biological variability into the quantification of differential AS by ranking inclusion differences based on how extreme they are with respect to the average transcript abundance they are associated with.

Yet, such tools, although of straightforward use to bioinformaticians, are not easily usable by and fully intelligible to non-computational scientists and/or lack interactive graphical frameworks allowing them to understand the statistics accommodating the two aforementioned sources of uncertainty, such that they can, for instance, critically evaluate discrepancies in consistency and reproducibility across differential AS analysis tools (12).

Therefore, we developed `betAS`, a user-friendly R package available via a web app that, from splice junction read counts obtained from `vast-tools` (16), `rMATS` (15), or `Whippet` (17)), allows intuitive analysis and visualisation of differential AS based on beta distributions. As mentioned, beta distributions are suitable to provide the estimated probability distributions of PSIs from the evidence for inclusion or skipping of alternative sequences given by read counts, such that each beta distribution has the observed PSI as its mean and is narrower when read coverage is higher. While the precision of AS estimates (PSIs) modelled with beta distributions is therefore proportional to the associated coverage and reflected on the significance of AS differences between samples, plotting the estimated beta distributions provides an intuitive graphical framework for evaluating the compromise between the uncertainty in individual sample estimates and the variability among replicates. Likewise, `betAS` thrives as a decision support system that helps the user to judge which AS changes are biologically relevant and robust, rather than using context-independent predefined cut-offs.

The betAS visual interface is designed to be used by any scientist, irrespectively of their prior knowledge of R, and simplifies the analysis of user-provided tables with AS quantifications, as well as the ranking of differentially spliced events by a significance metric that incorporates information on the precision of the relative transcript abundances and that is suitable for multiple-group comparisons. This feature, to our knowledge absent in the best established differential AS tools, is useful, for example, when studying the tissue- or differentiation stage-specific regulation of AS, as illustrated below in Results.

The betAS web app is available at https://compbio.imm.medicina.ulisboa.pt/betAS.

## MATERIALS AND METHODS

### Modelling inclusion levels from RNA-seq junction reads using the beta distribution

The betAS package and web app, developed in R (29), enable statistical differential alternative splicing analyses, and associated visualisation, from splice junction read count tables obtained from well-established AS quantification tools: vast-tools (16), rMATS (15), and Whippet (17).

AS can be quantified with RNA-seq. Transcripts in a biological sample are captured and fragmented for sequencing and the resulting reads are mapped to a reference of annotated gene structures, including information on the set of exons, introns and the splice junctions between these (i.e., exon-exon and exon-intron junctions) inferred for a representative set of each gene’s transcripts. An AS event is an alternative sequence, such as an exon, that is annotated as being included in some of the gene’s transcripts but not in others. In an RNA-seq sample, the fraction of mRNAs from a gene that contain a given alternative sequence is estimated from reads that, by mapping to exon-exon or exon-intron junctions, provide evidence for the inclusion or exclusion of that sequence across the cognate gene’s transcripts (Figure 1, left). Using these reads, proportion spliced-in (PSI) values (14) are calculated as the ratio between the number of reads that map to the junctions defining inclusion (inc) and the sum of these and reads that map to the junctions defining exclusion (exc) of that given alternative sequence (equation PSI, Figure 1, left).

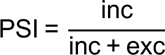

**Figure 1.**
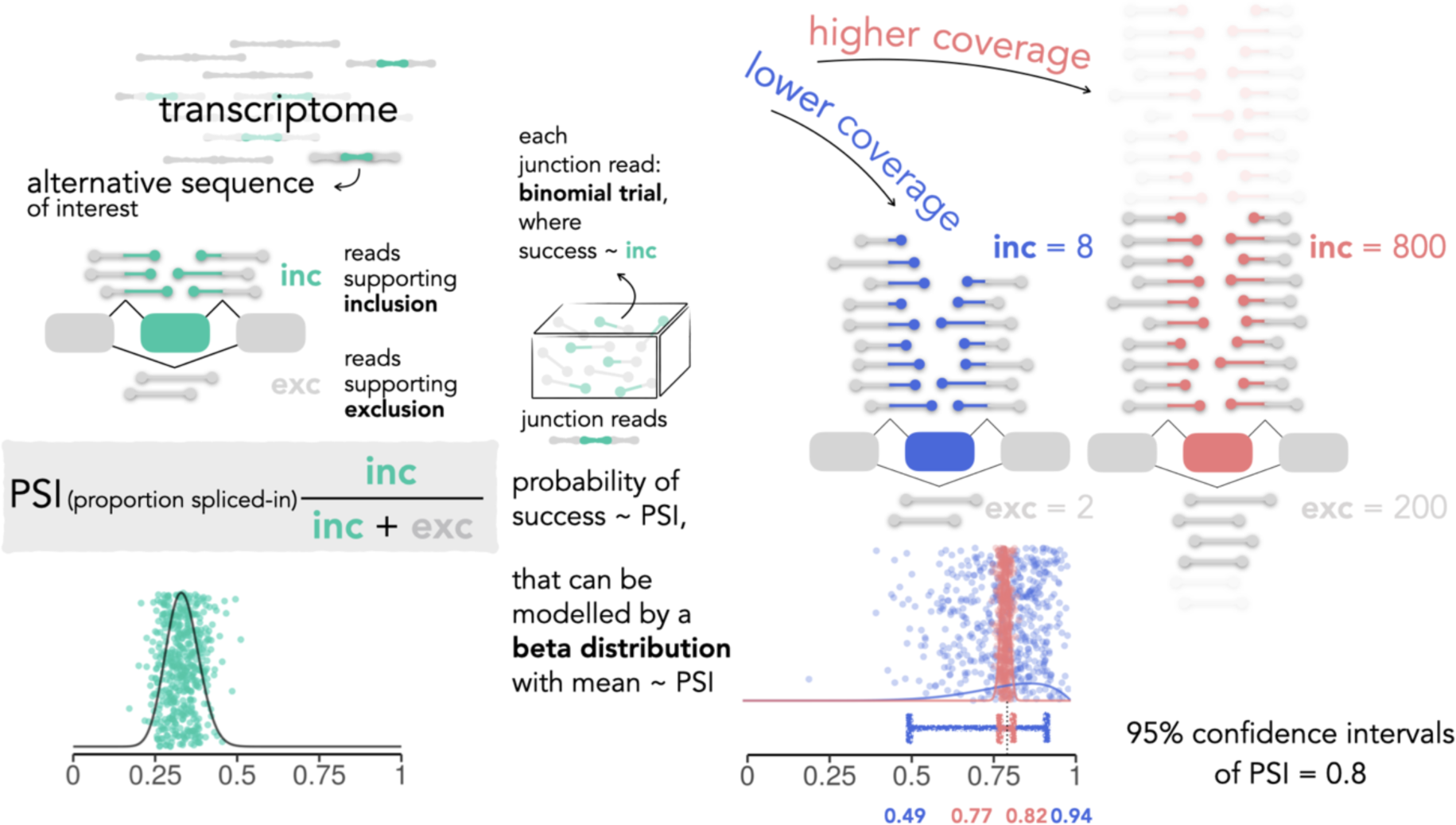
Beta distributions model PSI levels and associated confidence. Explanatory diagram of the estimation of the proportion spliced-in (PSI) with the beta distribution for an alternative transcript sequence (exon) of interest (green) from the RNA-seq junction read counts supporting the sequence’s inclusion (inc) or exclusion (exc). The ability of the beta distribution to incorporate the mean value and the confidence of the PSI is further illustrated for two discrepant scenarios of read coverage: lower, with 10 reads (blue), and higher, with 1000 reads (salmon). PSIs randomly generated from the respective beta distributions (500 coloured vertically “jittered” points per distribution and associated solid density lines) are dispersed around mean values, being less dispersed as coverage (i.e., confidence associated to supporting evidence) increases. Coloured 95% confidence intervals for a proportion’s test with p (in this case, PSI) equal to 0.8, with 10 (blue) or 1000 (salmon) trials. Coloured (green/blue/salmon) squircles: alternative exons; grey squircles: constitutive exons; junction read depictions coloured according to their exon coverage.

Most AS quantification tools use PSI values alone, that do not convey information on the number of reads used in the quantification, as the same PSI can be obtained with different levels of coverage (Figure 1, right). Beta distributions are useful in estimating and visualising the PSI’s precision, proportional to coverage (Figure 1 right and Supplementary Figure 1). The beta distribution is used to model phenomena with values constrained to [0,1], namely probabilities and proportions. The position and shape of the beta distribution are determined by two parameters that define the distribution’s mean and how narrow the distribution is (see *Supplementary Methods*).

In RNA-seq, coverage reflects the number of times an individual nucleotide is sequenced. Thus, when estimating PSI values, higher coverage implies higher confidence (i.e., precision) in that inclusion estimate. To model the PSI ratio and its precision for a particular AS event, the beta distribution’s shape parameters can conveniently incorporate information on the number of reads used in the quantification, inc and exc, such that the mean of the distribution is directly comparable to the PSI ratio (Figure 1), while for a given mean, the higher inc and exc are, the narrower the distribution is (Supplementary Figure 1).

Under the aforementioned motivation and improving on previous work by others (5, 15, 16, 19, 20), with a particular focus on the ideas underlying `vast-tools`’ *diff* module (30) and `Whippet` (17), the `betAS` package uses beta distributions to model alternative sequence inclusion levels (i.e., PSI values) from inc and exc reads associated with annotated AS events, and their graphical representation to facilitate the interpretation of differential AS between conditions.

### betAS approach for quantifying differential alternative splicing between two conditions

#### a) Estimate the effect size of splicing differences: ΔPSI

For each AS event, the modelling approach described above is used by `betAS` to estimate its inclusion levels per sample from the respective inc and exc read counts through the emission of random values (500 per sample by default) from a beta distribution with shape parameters inc and exc (Figure 2A). In cases where deemed to be enough high-coverage replicates to probe biological variability, it may be considered appropriate to weight the number of randomly emitted values proportionally to each sample’s coverage. `betAS` hosts this optional functionality, inactive by default. To profile differential AS between groups of samples, each AS event’s inclusion level in each replicate is first modelled using a beta distribution to reflect the individual estimate’s precision, proportional to the level of evidence (given by the read coverage) supporting that estimate (Figure 2A). A joint distribution encompassing each group’s replicate samples reflects both the estimation uncertainty in individual samples and the intra-group variability in PSI, therefore empirically modelling each group’s PSI and its precision, while providing a statistical framework for ΔPSI (i.e., difference in PSI between groups) quantification (Figure 2B).

**Figure 2.**
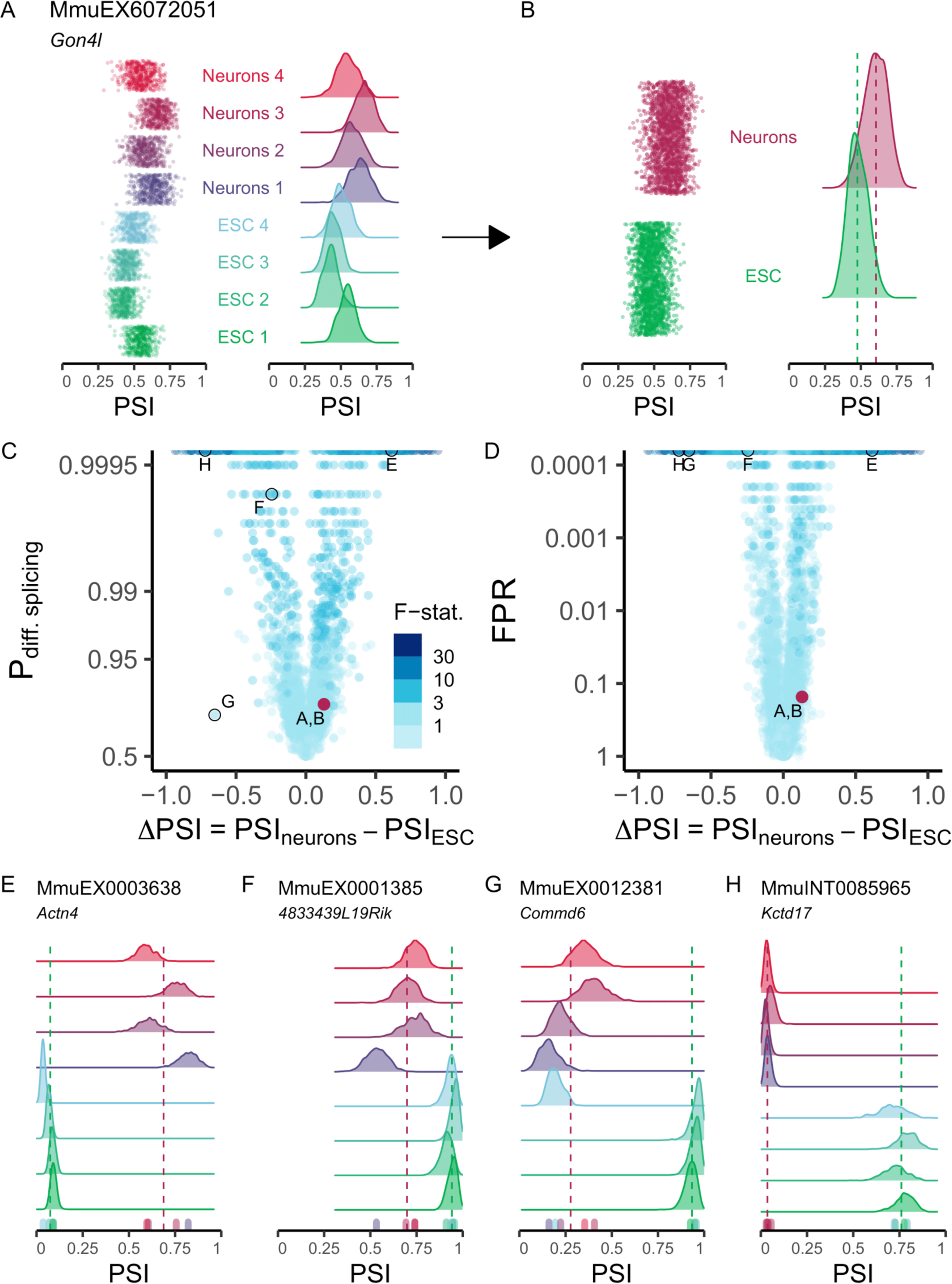
`betAS` pipeline enables intuitive visualisation of alternative splicing changes. (**A**, **B**) Beta distributions (vertical “jitter” and density plots of the emitted values) illustrating PSI dispersion for each sample (**A**) and per phenotypic group (**B**), with the vertical dashed lines indicating the median PSI for each group, PSI_neurons_ (dark red) and PSI_ESC_ (green). (**C**, **D**) Volcano plots illustrating the effect size (ΔPSI) as the difference between the median group PSIs and their significance assessed by the estimated probability of differential AS, based on the proportion of differences between ESC and Neurons beta distribution randomly emitted values that are > 0 (P_diff_) (**C**) and by the false positive rate (FPR) (**D**). Events are coloured by the F-statistic, i.e., the ratio of between-to within-group PSI variations (v. Materials and Methods). Both Y-axis scales are log-transformed to facilitate visualisation. Upper row of trimmed points represents P_diff_ > 0.9995 and FPR < 0.0001. (**E** to **H**) Beta distributions (density plots of the emitted values) and `vast-tools`’ PSIs (bottom coloured ticks) for selected events illustrative of different combinations of effect size and significance of AS differences, identified by `vast-tools`’ IDs (VAST-DB annotation for the mouse mm10 genome assembly): (**E**) MmuEX0003638 (gene *Actn4,* chr7:28895121-28895180), (**F**) MmuEX0001385 (gene *4833439L19Rik*, chr13:54564515-54564621), (**G**) MmuEX0012381 (gene *Commd6*, chr14:101640288-101640299) and (**H**) MmuINT0085965 (gene *Kctd17*, chr15:78436995-78438486). ESC: embryonic stem cells.

### b) Estimate the significance of splicing differences I: Pdiff

The first betAS approach to significance takes the two sets of random points per condition (Figure 2B) and calculates, for each AS event’s estimated ΔPSI, the proportion of differences between these that are greater than zero, which has the same interpretation as asking what proportion of beta distribution-emitted values for one condition are higher than those emitted for the other, thus reflecting the probability of differential AS, P_diff_ (equation Pdiff) of PSI_betAS_(**A**) being greater than PSI_betAS_ (**B**) (Figure 2C and Supplementary Figure 2).

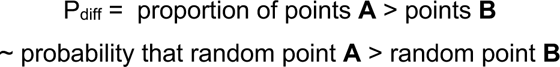

#### c) Estimate the significance of splicing differences II: FPR

`betAS` also allows the estimation of a false positive rate (FPR) for differential AS directly from the pipeline of random number generation. Following the emission of individual beta distributions per sample and to test against the null hypothesis that there is no difference in the PSI of a given AS event between two groups (ΔPSI = 0), i.e., that all PSI values in each sample of both conditions come from the same distribution, individual sample inc + exc values are used as the coverage of that event on that sample. As performed for the determination of a PSI per sample, random generation of numbers from a beta distribution is used to estimate the null distribution’s PSI and its precision. Thus, shape parameters determining each sample’s null beta distribution are given by random emission from a binomial distribution with number of trials, #trials = inc + exc and probability of success equal to the mean value of the PSI across all samples. This ensures that each sample’s null distribution inherits the PSI precision associated to their original coverage. Then, one number is randomly selected from each of the sample’s null distribution and, keeping the samples’ group assignment (i.e., which samples belong to each group), the ΔPSI between groups under the null hypothesis is calculated. The process is repeated many times (10 000 by default) and the FPR is the proportion of ΔPSI random simulations that are larger than (i.e., more extreme) or equal to the empirical ΔPSI (Figure 2D).

#### d) Estimate the ratio of between- and within-group variabilities of PSI: F-statistic

Capitalising on the incorporation of both the individual and group coverage-dependent PSI dispersion, `betAS` also enables an ANOVA-like analysis of variance, comparing inter- and intra-group variabilities. Thus, for each event, within is considered the set of differences between each pair of samples that are part of the same group and between the set of differences between each pair of groups. The ratio of the median absolute values of *between* and *within* therefore provides an “F-like” statistic (equation Fstat).

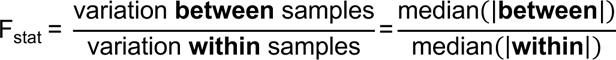

This metric provides a compromise between the effect size of AS differences and their significance, being therefore a suitable single metric for ranking AS events according to evidence for differential AS when both aspects are important (Figure 2C,D and Supplementary Figure 3).

### Simulation of RNA-seq splice junction read counts from empirically derived PSI and coverage values

Simulation of empirically derived PSI and coverage values was performed following a three-step approach (Figure 3A): first, GTEx (31) transcriptomic data (version 7) were used as a source of “ground truth” tissue-specific PSIs and “real” read coverage values from which junction read counts could be simulated; second, the biological variability of tissue-specific AS events was estimated by inspecting the PSI mean and variance of events with evidence for a unimodal PSI distribution (i.e., one typical PSI mode per tissue, thus assumed as tissue-specific) and finding the variance/mean relationship of low-variance events (see *Supplementary Methods*); third, for the same set of representative unimodal AS events, junction read counts were simulated for a given number (N) of replicates by incorporating the biological variability estimated in step 2 and including technical variability (i.e., variation amongst replicates), both inspired in the “ground truth” PSIs and empirical coverages mentioned in step 1.

**Figure 3.**
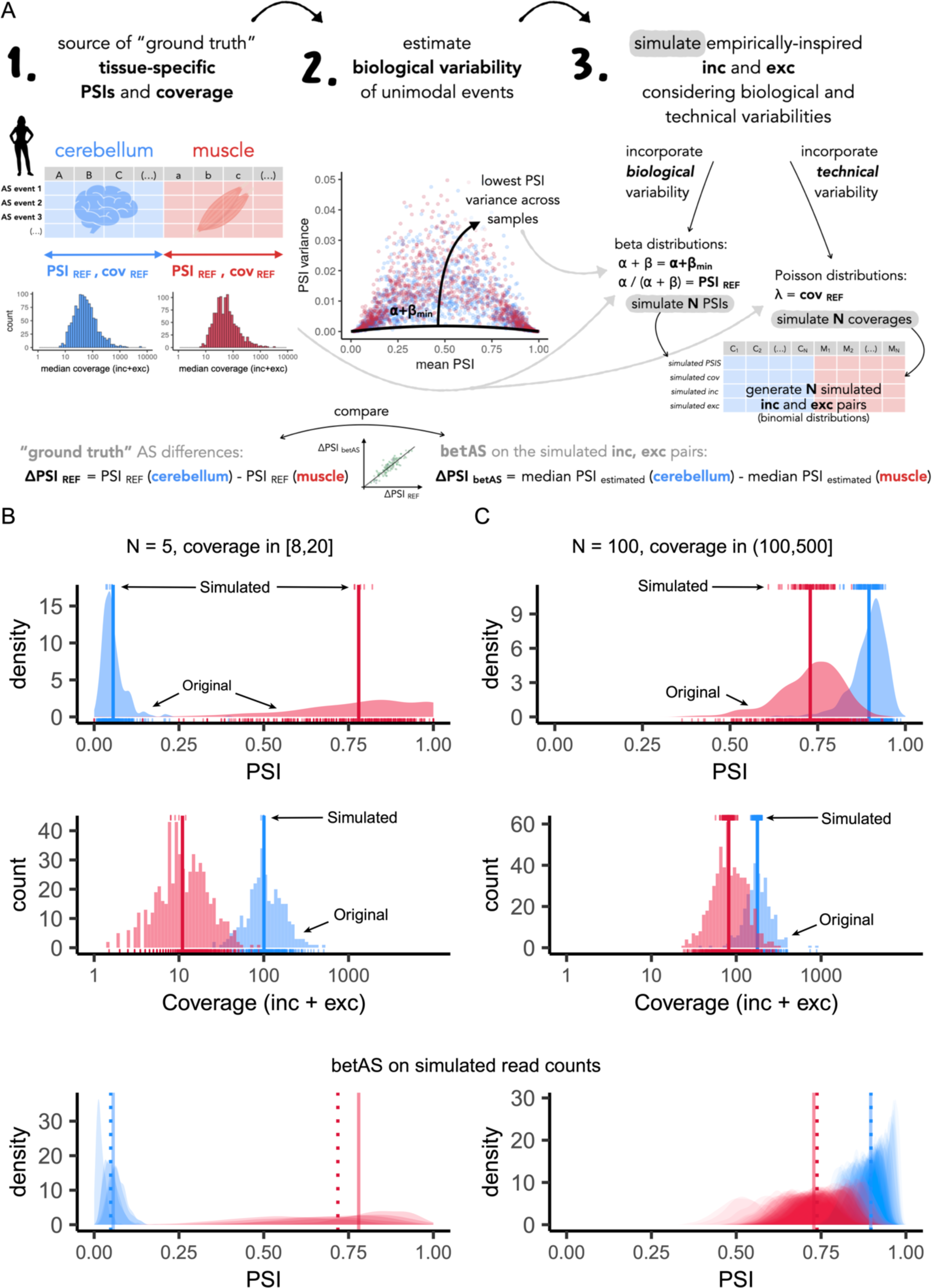
`betAS` accuracy in measuring PSI differences from simulated read counts. (**A**) Explanatory diagram of the implemented approach to simulate empirically-inspired RNA-seq junction read counts. Reference PSI (PSI_REF_) and coverage (cov_REF_) values for 563 muscle (red) and 309 cerebellum (blue) samples in GTEx version 7 are used to generate N (number of replicates) pairs of randomly emitted low-variance PSI and coverage values per event, used to simulate RNA-seq junction read counts on which `betAS` accuracy in measuring PSI differences can be evaluated. (**B** and **C**) Examples of the simulation approach applied over a low coverage, cov_REF_ in [8,20] junction read counts, and low number of replicates, N = 5, event (**B**) and over a high coverage, cov_REF_ in [100,500] junction read counts, high number of replicates, N = 100, event (**C**). Top panels: density and bottom rug plot showing the original distribution of PSI values for all samples; top rug plot showing the randomly emitted (“simulated”) low-variance PSIs. Vertical lines indicate PSI_REF_ values for muscle and cerebellum. Middle panels: histogram and bottom rug plot showing the original distribution of coverage values for all samples; top rug plot showing the randomly emitted coverage values. Vertical lines indicate cov_REF_ values for muscle and cerebellum. Bottom panels, “betAS on simulated read counts”: density plots representing PSI estimation as done in `betAS`, with values randomly emitted from a beta distribution with the shape associated with the simulated junction read counts per sample. Solid vertical lines indicate PSI_REF_ values for muscle and cerebellum, while dashed vertical lines indicate the median PSI values estimated by `betAS` for muscle and cerebellum.

In detail, functions from `psichomics` (32) were used to load sample, subject and junction information for 563 muscle and 309 cerebellum samples (Figure 3A, left). PSIs were quantified for 38 896 exon skipping events using the *quantifySplicing* function (minReads = 0). The bimodality coefficient can be calculated based on the shape of a distribution and has been proposed as suggestive of bimodality if > 5/9 (33). Thus, in order to select a set of exon skipping events with tissue-specific unimodality amongst the 14 039 events with valid PSIs for all samples, only events with a PSI bimodality coefficient (calculated with *modes* R package, available as an archive at https://cran.r-project.org/src/contrib/Archive/modes) per tissue < 3/9 in Muscle and Cerebellum were considered (Supplementary Figure 4A).

In order to empirically estimate the minimum biological variability of real tissue-specific events (Figure 3A, middle), the relationship between the PSI variance and mean PSI per tissue for the subset of 1 143 tissue-unimodal events (Supplementary Figure 4B-D) was assessed to identify the subset of such events that were associated with the lowest variance, i.e., the tissue-unimodal events with the least observed biological noise (see *Supplementary Methods*).

Thus, for each tissue-unimodal AS event used as “biological inspiration”, `betAS` can simulate a given number (N) of replicate PSIs by the following procedure (Figure 3A, right, and Supplementary Figure 4E; see *Supplementary Methods*): first, considering all samples, use the mean of the distribution of GTEx PSI values (“Original PSIs”, Supplementary Figure 4E top left) as the reference PSI (PSI_REF_) of that event per tissue and simulate, for each tissue, N PSI values from a beta distribution with shape parameters such that α + β = 138.5042 (empirically inferred, as described in Figure 3A and Supplementary Figure 4D; see *Supplementary Methods*) and α/(α + β) = PSI_REF_ (“Simulated PSIs”, Supplementary Figure 4E middle left); using the median of the GTEx coverages (cov_REF_) associated with the original PSIs (“Original coverage”, Supplementary Figure 4E top right), sample N instances of cov from a Poisson distribution with λ = cov_REF_ (“Simulated coverage”, Supplementary Figure 4E middle right). Sampling PSI and coverage values separately ensures these likely come from different samples of the same tissue, even though they are associated with the same event. Finally, by pairing simulated PSI and coverage values, the *rbinom* function is used to generate, from a binomial distribution, inc and exc read pairs per simulated replicate, to which `betAS` is applied (Supplementary Figure 4E, “betAS”, bottom left). Differences in simulated PSI values for different scenarios of N (5, 10, 50 and 100) between tissues (ΔPSI) can be obtained and compared (Figure 3A, bottom) with respective reference differences:

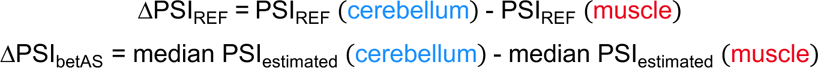

### betAS applied to comparisons between multiple groups

The F-statistic can inherently be used in testing differential AS between more than two groups and the `betAS` pipeline extends the approach described above with that purpose. The ratio of the median absolute values of between (i.e., distances between random points from samples in one group and points in the other groups) and within is again translated into the “F-like” statistic for the multiple-group comparison. Likewise, the application of the P_diff_ between two groups can be extended to encompass the comparison between multiple groups (see *Supplementary Methods*).

Moreover, for the particular case of experimental designs with groups following a sequential relationship (e.g., timecourse), `betAS` defines sequential between as the between-group differences for any sequential pair of groups. Comparing the sign of the differences between each group and its predecessor, `betAS` can infer if inclusion levels are increasing or decreasing along the sequence (i.e., if there is a monotonic trend), as well as estimate a monotonicity coefficient across the complete course (see *Supplementary Methods*).

While there is no technical restriction on the maximum number of groups to be compared, the practical limitation lies in the visualisation and interpretation of results. As the number of groups increases, interpretability becomes more challenging. Metrics can aid in prioritizing events, but visual inspection remains crucial for interpretation.

### betAS visual interface

Our pipeline for differential AS analyses is accessible online at https://compbio.imm.medicina.ulisboa.pt/betAS. `betAS` web app was designed to guide users that are not familiar with programming in analysing their own differential AS experiments. `betAS` web app can take as input a tab-delimited text file including inclusion level quantifications (PSI) together with both the inclusion and exclusion junction read counts supporting each PSI.

The app provides a launching first tab, entitled “Import inclusion levels”, that allows loading a table with AS quantifications or, by default, proceeding with an example table obtained from an RNA-seq time-series experiment on the development of seven major human organs (34). At this stage, and following filtering out events with very low coverage (see *Supplementary Methods*), users can select the AS event types (e.g., exon skipping, intron retention, etc.) to analyse, as well as the PSI range within which to consider quantifications for analysis, while they are visually informed on the features of the population of selected events through dynamic plots and tables.

The “Group definition” tab allows users to explicitly convey the experimental design underlying their questions (sample annotation). Groups of samples can be manually generated by introducing their names, identifying their samples and assigning colours. Alternatively, `betAS` can automatically screen for similarities in the samples’ names (typically the column names in the input PSI table) under the “Automatic group(s)” option, or, in the case of the default example table, use a provided feature to automatically generate groups, via the “Feature-associated group(s)” button. Groups can be deleted and created anytime by the user.

The web app supports both pairwise and multiple-group differential AS analyses. Following group definition, running differential AS analysis between groups with `betAS` is straightforward, involving choosing the y-axis significance metric to consider for the volcano plots and, in the case of 2 groups, the groups of interest for the comparison. Owing to the stochastic nature of beta distribution random number generation in `betAS`, each run may produce distinct outcomes. This variability poses a challenge, especially for events near significance thresholds. To address this, we introduced the option to set a seed when executing `betAS`, offering a practical means for reproducibility. A 5-minute video tutorial on the app, to guide the user over its sections and features, is available at https://compbio.imm.medicina.ulisboa.pt/betAS/tutorial.

The betAS package, whose source code is available at https://github.com/DiseaseTranscriptomicsLab/betAS, includes functions to generate beta distributions, calculate metrics of differential AS, generate volcano plots for the global analysis of all events, as well as AS event-specific plots illustrating differential AS (such as density plots summarising the distributions of both coverage and inclusion levels across samples).

## RESULTS

### Beta distributions model PSI levels and associated confidence

Profiling alternative splicing from an RNA-seq-derived transcriptome relies on sampling the mRNA molecules present in the studied tissue at a given moment. RNA-seq reads that map to annotated exon-exon or exon-intron junctions are typically used as supporting evidence for either inclusion or exclusion of alternative sequences in transcripts. The relative inclusion level of a known alternative sequence in a biological sample can be conveniently modelled by the beta distribution, with shape parameters given by the number of inclusion and exclusion reads (Figure 1 and Supplementary Figure 1; see Materials and Methods) (15, 17, 19, 20, 30), with the precision of the inclusion level estimate being therefore proportional to the number of supporting reads.

As shown in Figure 1, for a particular AS event the PSI quantification alone provides no information on the total number of supporting RNA-seq reads but only on the proportion of them providing evidence for inclusion. The two depicted AS events have contrasting number of reads supporting inclusion/exclusion (8/2 for the lower-coverage example in blue, 800/200 for the higher-coverage case in salmon) but show the same PSI value. However, the stronger evidence provided by higher coverage (salmon) confers more confidence to its PSI estimate than to that of a case with lower coverage (blue), as illustrated by their confidence intervals (Figure 1).

### betAS pipeline enables intuitive visualisation of the magnitude and significance of alternative splicing changes

To provide a decision-support tool for the analysis of AS differences between conditions, allowing subsequent interpretation of their biological implications, betAS implements intuitive visuals that incorporate the estimated inclusion levels and their precision.

AS is more prevalent and more conserved, across vertebrates, in nervous system tissues (35, 36). Moreover, neuronal-specific RNA-binding proteins that regulate AS in neurons contribute to the functional complexity of neurodevelopmental processes (37, 38). Given this relevance of AS in neuronal specification, as a case study, `betAS` was applied to the analysis of AS changes during the transition from pluripotency to mature neuronal function in mouse by comparing the transcriptomes of samples in the extremes of the publicly available murine neuronal differentiation timeline (39): embryonic stem cells (ESCs) and mature neurons (28 days *in vitro*).

Volcano plots illustrating the effect size/significance relationship for PSI differences across AS events is shown in Figure 2C-D. In this example, the selection of events differentially spliced between the two conditions aims to maximise both the probability of their PSI values being greater in one of the conditions and the actual difference in PSI between the two conditions. Thus, higher values of P_diff_ and lower values of FPR highlight more robust separations between the estimated distributions of PSI (Supplementary Figures 2 and 3), while greater effect sizes (ΔPSI) suggest stronger impacts on sequence inclusion of the biological differences between conditions.

`betAS` allows the interpretation of ΔPSI values associated with different PSI/coverage scenarios. For instance, while the selected *Gon4l* exon (Figure 2A-B) is barely more included in neurons than in ESCs, the volcano plot illustrates some stronger and more robust AS changes (Figure 2C-D), with several AS events showing switch-like behaviours, as seen by extreme differences (i.e., close to absolute values of 1) in PSI between neurons and ESCs (Figure 2C-D). Some of these AS changes are illustrated in Figure 2E-H, where the visualisation of the beta distributions that model sample-wise inclusion levels elucidates on the differences at the PSI levels and their precision across samples. Amongst them, the two most extreme examples in significance and effect size, with absolute ΔPSI values greater than 0.50, are exon MmuEX0003638 in gene *Actn4*, rarely included (PSI ≈ 0) in ESC but typically included (PSI ≈ 0.70) in mature neurons (Figure 2E) and in neuronal tissues (https://vastdb.crg.eu/event/MmuEX0003638@mm10), and intron MmuINT0085965 in *Kctd17*, that shows high retention levels in ESC (PSI ≈ 0.75) and is spliced out (PSI ≈ 0) in neurons (Figure 2H). In both cases there are coverage differences (associated with differences in gene expression) between the two conditions, reflected by the different widths of the respective beta distributions (Figure 2E,H). Visualisation of inclusion levels through density plots of the respective beta distributions together with the usage of associated significance metrics such as the P_diff_ also allow the user to spot outlier replicates, as for exon MmuEX0012381 in *Commd6* (Figure 2G), nearly constitutively included (PSI ≈ 1) in all ESC samples except one (PSI ≈ 0.2). Although showing an average ΔPSI similar to that of the intron retention event in Figure 2H, its P_diff_ significance is penalised due to that inconsistency between replicates (Figure 2C).

### Assessment of betAS ability to estimate differential AS on simulated RNA-seq splice junction read counts from empirically derived PSI and coverage values

In order to benchmark `betAS`‘ accuracy in quantifying differential AS, it should be tested on simulated RNA-seq junction read counts, such that the ability of the ΔPSI values estimated with `betAS` from the simulated reads could be compared with “ground truth”, reference ΔPSI values. Therefore, the biological and technical variability in PSI and read count distributions and their association with the sequencing sampling, together with the features of beta and Poisson distributions, were explored to simulate RNA-seq spliced junction read counts inspired in a real biological dataset (see *Supplementary Methods*).

Large-scale AS quantifications available from GTEx (31) were used to simulate “biology-inspired” PSI values (Figure 3A) (see Materials and Methods). To simulate PSIs reflecting the natural inclusion patterns of highly regulated AS events, the focus was on a) tissues with prevalent and functionally relevant AS (muscle and brain); and b) AS events with typical inclusion levels that are lowly variable in samples of the same tissue, properties that find a proxy in the mathematical concept of unimodality of a distribution. For each representative tissue-unimodal AS event, junction read counts were simulated for a set of N replicates from PSI/coverage pairs.

Differences in inclusion between cerebellum and muscle estimated with `betAS` from the simulated junction read counts (ΔPSI_estimated_) were compared with those found for the reference tissue PSIs per event in GTEx (ΔPSI_REF_) (Figure 3A and Supplementary Figures 4 and 5). As expected, `betAS` accuracy in estimating the “real” ΔPSI increases with the simulated number of replicates (N) and coverage (Supplementary Figure 5), as illustrated by the selected illustrative examples in Figure 3B,C. Moreover, discrepancies found between ΔPSI_simulation_ and ΔPSI_REF_ are more pronounced for median PSI values in the alternative region in the middle of the [0,1] range, i.e., around 0.5 (in green in Supplementary Figure 5). This is expected due to the “scaling effect” (40), i.e., alternatively spliced sequences with intermediate PSI levels are more likely to undergo larger ΔPSIs as a response to perturbations than those nearly-constitutively included or excluded.

In any case, even considering low replication level and coverage (Figure 3B and Supplementary Figure 5), betAS accurately estimates real ΔPSI values, being therefore a suitable tool for differential AS analysis.

Differential AS significance metrics P_diff_ and FPR introduced by `betAS` will naturally depend on coverage and ΔPSI values. In a simulated typical scenario of differential AS analysis between two groups of three biological replicates each, an arbitrary P_diff_ cut-off of 0.95 will allow significant detection of events with |ΔPSI| values higher than ∼0.3, ∼0.2, and ∼0.1 for coverages of 10, 100, and 1000 reads, respectively (Supplementary Figure 6A). Similarly, an arbitrary FPR cut-off of 0.05 will allow significant detection of events with |ΔPSI| values higher than ∼0.2, ∼0.15, and ∼0.05 for those coverages (Supplementary Figure 6B). Despite the proportionality between P_diff_ and FPR (Supplementary Figures 3A and 6C), that discrepancy between the two metrics in the minimum effect size detectable with a typical significance cut-off of 5% is explained by them representing different meanings of significance (see Materials and Methods), therefore providing complementary information to support users’ decisions on differential AS. The observed variability in the minimum |ΔPSI| values detectable for fixed P_diff_/FPR cut-off and coverage is due to the aforementioned “scaling effect”, i.e., the precision of quantification of ΔPSI values depends on the actual range of PSI valuesinvolved.

### Visual comparison of betAS with other tools for differential AS analyses

The probability of differential AS P_diff_ and FPR introduced by `betAS` were compared to significance metrics from other well-established methods for differential splicing analysis with matching AS event annotation (see *Supplementary Methods*): `SUPPA2` (18), `rMATS` (15), and `Whippet` (17) (Supplementary Figure 7). As in `betAS` applied on junction read counts, both `SUPPA2`’s and `rMATS`’s effect size and significance metrics allow the visualisation of the differential AS results through a volcano plot. However, neither `SUPPA2` or `rMATS` are oriented towards allowing visual inspection of how each replicate contributes to its estimated group PSI and that contribution reflects on the differential AS estimates, something particularly relevant when dealing with few samples. Thus, using `betAS` visualisation infrastructure on differential AS metrics obtained with those tools contributes to make their metrics directly comparable and interpretable. Results of differential AS in mouse neuronal development obtained with each tool were compared for the same set of alternative exons.

`SUPPA2`’s differential AS approach relies on monitoring the uncertainty in PSI estimates as a proxy of biological variability. Since low coverage is associated with higher variability, `SUPPA2`’s assumption is that the significance of each observed ΔPSI between conditions depends on where in the distribution of coverage/uncertainty that difference is (18). Thus, the same ΔPSI can be considered significant if falling in a low uncertainty (high coverage) region while that might not be the case if it falls in the high uncertainty (low coverage) region. Likewise, each ΔPSI obtained is evaluated as more or less extreme for its cognate transcript average coverage level. `SUPPA2`’s estimates of significance of AS differences are supported by showing ΔPSI values between replicates as a function of the average transcript abundance (Supplementary Figure 8D and 8E).

The significance cut-off suggested by `SUPPA2` (see *Supplementary Methods*) and a P_diff_ > 0.95 were considered to study differences in differential AS calls identified by `betAS` and `SUPPA2`. Inspection of events that are selected as differentially spliced by both, one or none of the approaches, i.e., cases in each “quadrant” of the scatter plot comparing the respective significance metrics (Figure 4A), elucidates on the estimates underlying the significance calls for AS differences made by `SUPPA2` and `betAS`. For instance, exon MmuEX0051014 in *Usp4* (Figure 4F), considered significantly differentially spliced by `betAS` based on an estimated P_diff_ > 0.95, has PSI estimates per sample such that, even showing a small ΔPSI, their group probability distributions are separated enough between ESC and neurons to be considered different under the P_diff_ cut-off considered. For `SUPPA2`, however, such a small ΔPSI is not sufficient to be considered significant at that average coverage (Supplementary Figure 8D). On the other hand, exon MmuEX0024526 in *Isca2* (Figure 4G) is considered significantly differentially spliced by `SUPPA2`, as its ΔPSI is amongst the most extreme in its range of coverage (Supplementary Figure 8D), but not by `betAS` based on P_diff_, due to the relatively high inter-replicate variability illustrated by the overlap of PSI distributions between ESC and neurons (Figure 4G, bottom).

**Figure 4.**
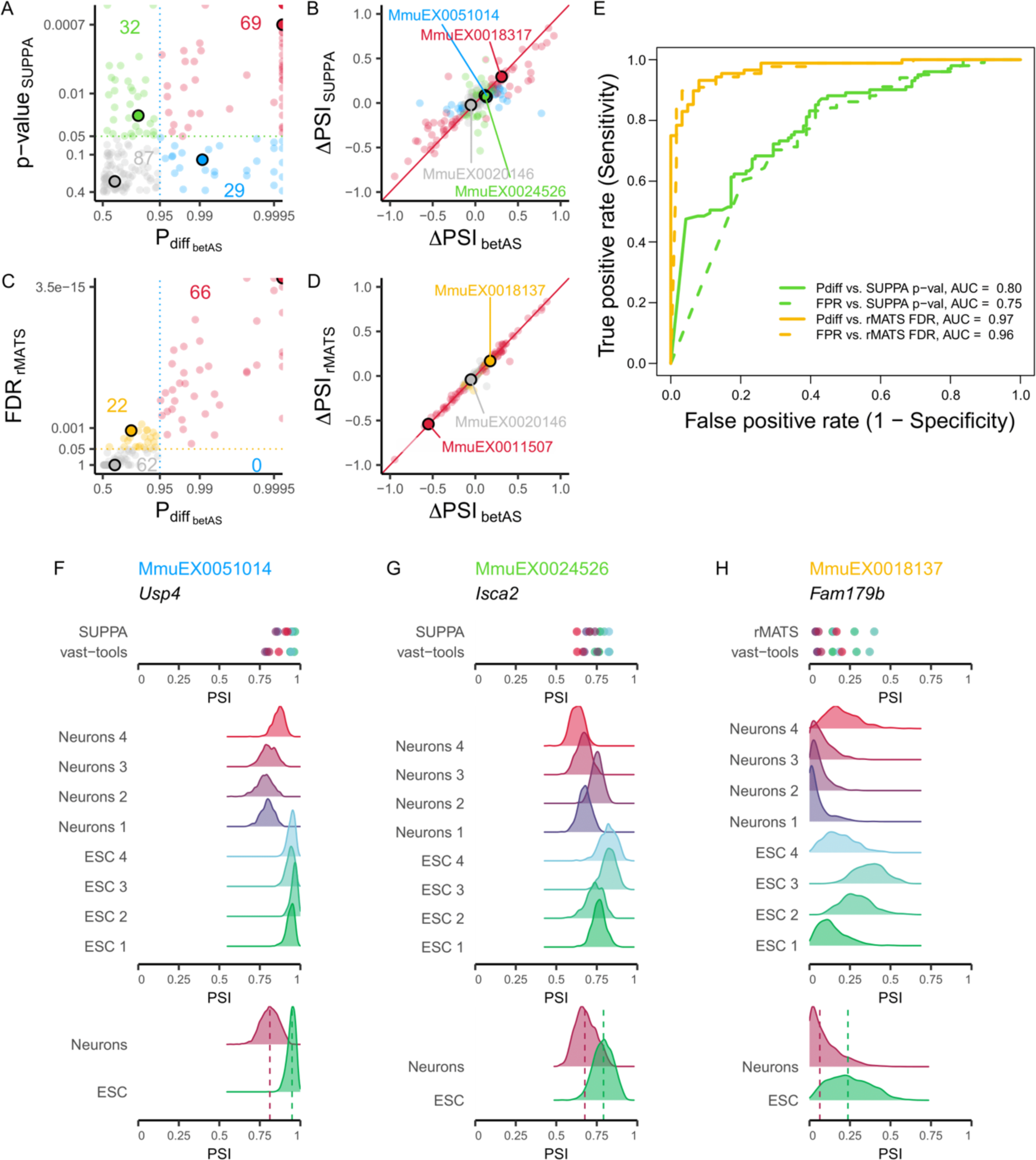
Comparison of significance of differential AS between `betAS`, `SUPPA` and `rMATS`. (**A**) Scatterplot comparing `betAS`’ and `SUPPA`’s differential AS significance metrics. `betAS`’ estimated probability of differential AS, based on the proportion of differences between the beta distributed randomly emitted values per group that are > 0 (P_diff_) and `SUPPA`’s p-value, with dashed lines defining quadrants indicating the significance cut-offs considered (P_diff_ > 0.95 for `betAS`, p-value < 0.05 for `SUPPA`). Points are coloured based on the differential AS calls by both tools: events considered differentially spliced by both `betAS` and `SUPPA` (red), by `betAS` (blue) or `SUPPA` (green) alone or by none of the tools (grey). Selected example events for each quadrant as larger outlined dots. (**B**) Scatterplot comparing `betAS`’ and `SUPPA`’s differential AS effect size (ΔPSI). Red diagonal solid line indicates identity: ΔPSI_SUPPA_ = ΔPSI_betAS_. (**C**) Scatterplot comparing `betAS`’ and `rMATS`’s differential AS significance metrics. `betAS`’ P_diff_ and `rMATS`’s FDR, with dashed lines defining quadrants indicating the significance cut-offs considered (P_diff_ > 0.95 for `betAS`, FDR < 0.05 for `rMATS`). Points are coloured based on the differential AS calls by both tools: events considered differentially spliced by both `betAS` and `rMATS` (red), by `betAS` (blue) or `rMATS` (yellow) alone or by none of the tools (grey). Selected example events for each quadrant as larger outlined dots. (**D**) Scatterplot comparing betAS’ and `rMATS`’s differential AS effect size (ΔPSI). Red diagonal solid line indicates identity: ΔPSI_rMATS_ = ΔPSI_betAS_. (**E**) Receiving Operating Characteristic (ROC) curves `betAS`’ differential AS calls with P_diff_ (solid lines) and FPR (dashed lines), considering as ground truth the differential calls from `SUPPA` (yellow, p-value < 0.05) or `rMATS` (green, FDR < 0.05). (**F** to **H**) `SUPPA` and `vast-tools`’ PSIs (top) and densities of emitted beta distributed values for individual samples (middle) and their merging per sample group (bottom, with dashed lines signing median values) for selected events illustrative of different combinations of effect size and significance of AS differences, identified by `vast-tools`’ IDs (VAST-DB annotation for the mouse mm10 genome assembly): (**F**) MmuEX0051014 (gene *Usp4,* chr9:108388277-108388380), (**G**) MmuEX0024526 (gene *Isca2*, chr12:84773793-84773908), (**H**) MmuEX0018137 (gene *Fam179b*, chr12:64990941-64991090). ESC: embryonic stem cells.

`rMATS`’s differential AS statistical approach relies on modelling inclusion levels by both accounting for the PSI uncertainty in individual replicates, dependent on the total number of supporting RNA-seq reads, and the (biological) variability of PSI values across replicates (15). `betAS`’ ΔPSI values are generally more consistent with those obtained by `rMATS` (Figure 4D) than with those obtained by `SUPPA2` (Figure 4B), reflecting the consistency between `vast-tools` and `rMATS` PSI quantification. Although rMATS’ FDR is more sensitive than `betAS`’ P_diff_ at the selected cut-off (Figure 4C,E and Supplementary Figure 9A-C), the significance assessment by `betAS` and `rMATS` is also more consistent than between `betAS` and `SUPPA2`. One example illustrative of the difference in sensitivity is exon MmuEX0018137 in *Fam179b*, whose lower inclusion in mature neurons compared to ESC (Figure 4H) is considered significant by `rMATS`, but `betAS` P_diff_ is < 0.95. Visual inspection of the inclusion levels summarised with beta distributions again uncovers high inter-replicate variability illustrated by the overlap of PSI distributions between ESCs and neurons, that penalises the P_diff_ (Figure 4H), in contrast with the MmuEX0011507 exon in *Clasp1*, considered significantly differentially spliced between ESCs and neurons by both tools (Supplementary Figure 9F), consistently with its known increased inclusion levels in neural tissues (https://vastdb.crg.eu/event/MmuEX0011507@mm10). Similar observations are made when choosing FPR as the significance metric for `betAS` (Supplementary Figures 8C and 9C).

The comparison of differential AS between betAS and Whippet (Supplementary Figure 10) leads to similar observations as that between betAS and rMATS.

### betAS applied to differential AS between multiple groups

The differential AS approach implemented by `betAS` can be applied to multiple (i.e., more than two) groups in a novel ANOVA-inspired way that extends the P_diff_ definition to the comparison of the differences between samples belonging to different biological conditions to those found between replicates. Statistics for multi-group differential AS are, to our knowledge, absent from the aforementioned established tools and therefore a unique feature of `betAS`, applicable to the study of splicing in a variety of biological contexts.

To illustrate this, we applied multiple-group `betAS` to the analysis of AS differences in a subset of human transcriptomes of forebrain, hindbrain, heart, kidney, liver and testis (34) (Figure 5). By comparing the medians of differences “between” and “within” groups, both through their difference and their ratio (Figure 5A and B), while visually inspecting their distributions (Figure 5C-F, panels on the left), tissue-specific AS can be identified (Figure 5C-F). While the increased number of samples renders the individual beta distributions less relevant in terms of visualisation, the probability that |between|>|within| is, in this case, useful to identify AS events with differences between tissues. The selected exon HsaEX0021044 in *DST*, for instance, is more commonly skipped in kidney, liver and testis, whether its inclusion is higher and more variable in brain and heart tissues (Figure 5C). On the contrary, exon HsaEX0051256 in *PTS* is more excluded in brain and heart but less so in kidney, liver and testis (Figure 5E).

**Figure 5.**
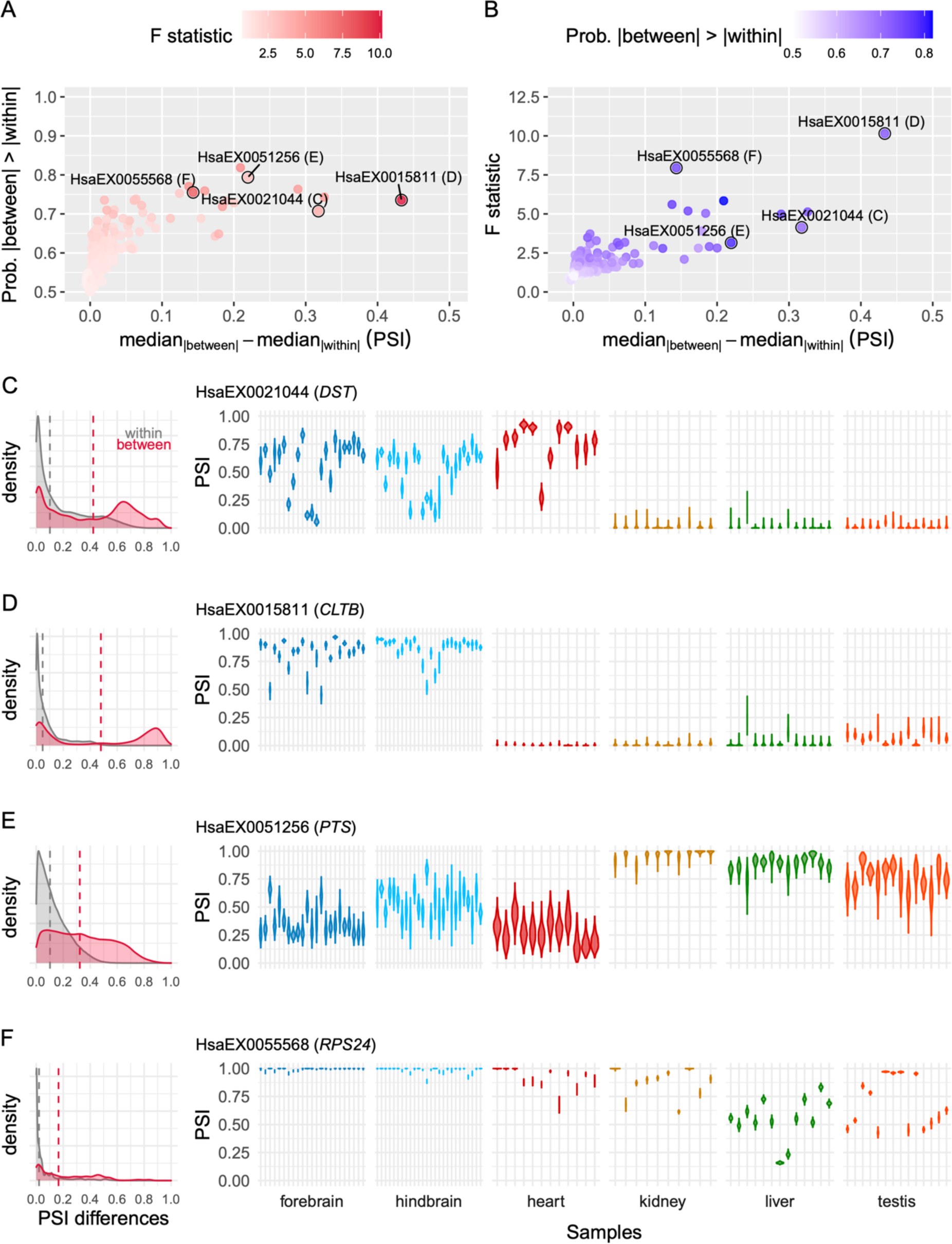
Beta distributions model inclusion level differences across multiple groups. (**A**) Scatterplot comparing `betAS`’ difference of the median of absolute differences between and within groups and the estimated probability that absolute differences between groups are greater than those within groups. Points are coloured based on estimated F statistic as the median of absolute differences between divided by the median of absolute differences within groups. Highlighted points are associated with the selected example AS events in panels C to F. (**B**) Scatterplot comparing `betAS`’ difference of the median of absolute differences between and within groups and the estimated F statistic as the median of absolute differences between divided by the median of absolute differences within groups. Points are coloured based on the estimated probability that absolute differences between groups are greater than those within groups. Highlighted points are associated with the selected examples in panels C to F. (**C** to **F**) Left: density plots of the absolute differences between (red) and within (grey), with dashed vertical lines indicative of the median values of each distribution. Right: Beta distributions (violin plots of the emitted values) for selected events, identified by `vast-tools`’ IDs (VAST-DB annotation for the human hg19 genome assembly): (**C**) HsaEX0021044 (gene *DST,* chr6:56329483-56329554), (**D**) HsaEX0015811 (gene *CLTB*, chr5:175823480-175823533), (**E**) HsaEX0051256 (gene *PTS*, chr11:112100931-112100953), (**F**) HsaEX0055568 (gene *RPS24*, chr10:79799962-79799983).

In order to explore the tissue specificity of AS events or more general questions about AS differences across groups, `betAS` allows flagging the more variable events across groups (Figure 5A and 5B), while inspection of the underlying beta distributions (Figure 5C-F) can inform on which conditions are driving the differences. Moreover, applying the F statistic to differential AS quantifications allows ranking events based on how robust the differences between groups are when compared to the differences between samples in the same phenotypical group (Figure 5A and 5B, Supplementary Figure 3).

`betAS` can also be used to study differential AS across groups with a sequential association, e.g., the mouse neuronal differentiation timeline dataset (39) (Figure 6). In this case, differences between sequential groups inform on the “steps”, or the monotonicity of the variation of inclusion levels (Figure 6A and B). Thus, “sequential” differences between groups together with their number of modes may clarify on the monotonicity of transitions in inclusion levels, such as those illustrated by event MmuEX0045617 in *Stxbp1*, whose PSI decreases from ESC to nearly full skipping around 0 *days in vitro* and then progressively increases as neurons differentiate (Figure 6C). The number of modes (or “step heights”) of the “sequential” differences and of the “generalised” differences (the median of differences found between groups in the extremes of the timeline) can be useful to interpret the progression of PSI along a sequence. For instance, the inclusion of exon MmuEX0024060 in *Ilf3* (Figure 6E) decreases monotonically (from 0 until 16 *days in vitro*) with multiple “steps” of the same size (PSI difference), thus showing one sequential mode that is different from 0. Exon MmuEX0016666 in *Elavl2* (Figure 6F) shows a PSI increasing in the early differentiation timepoints (from −4 to 0 and then to 1 *days in vitro*), consistent with its function in safeguarding loss of neuronal ELAV function in *Drosophila* (41) with a PSI “step” that is represented by the positive sequential mode identified.

**Figure 6.**
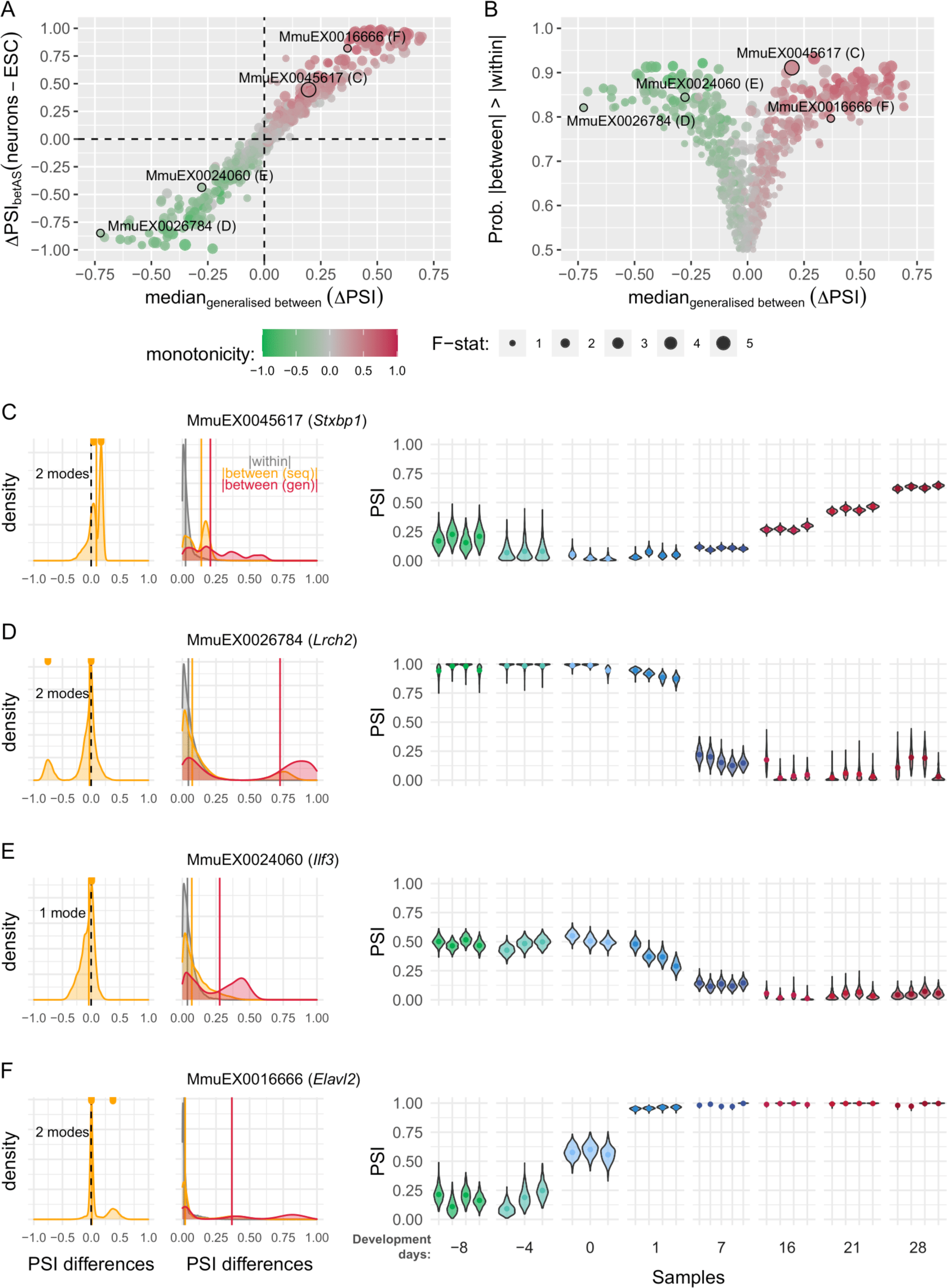
Beta distributions model inclusion level differences across sequential groups. (**A**) Scatterplot comparing `betAS`’ median “generalised” differences between groups, i.e., the median of differences found between groups and the effect size (ΔPSI_betAS_) between the extremes of the timeline (development days 28 – neurons - to −8 days - ESC). Points are coloured based on estimated monotonicity coefficient (see Supplementary Methods) and shaped based on F statistic as the median of absolute differences between divided by the median of absolute differences within groups. Highlighted points are associated with the selected example AS events in panels C to F. (**B**) Volcano plot comparing `betAS`’ median “generalised” differences between groups and the estimated probability that absolute differences between groups are greater than those within groups. Points are coloured based on estimated F statistic as the median of absolute differences between divided by the median of absolute differences within groups. Highlighted points are associated with the selected examples in panels C to F. (**C** to **F**) Left: density plots of the PSI differences between sequential groups (orange) in the timeline with rug plots (top) indicating the identified modes, with dashed vertical black line indicative of 0. Middle: density plots of the absolute “generalised” (red), sequential (orange) and within (grey) groups differences in PSIs, with vertical lines indicative of the median values of each distribution. Right: beta distributions (violin plots of the emitted values, with point indicating the mean of each distribution) for selected events, identified by `vast-tools`’ IDs (VAST-DB annotation for the mouse mm10 genome assembly): (**C**) MmuEX0045617 (gene *Stxbp1,* chr2:32794588-32794713), (**D**) MmuEX0026784 (gene *Lrch2*, chrX:147484754-147484804), (**E**) MmuEX0024060 (gene *Ilf3*, chr9:21388112-21388150), (**F**) MmuEX0016666 (gene *Elavl2*, chr4:91254141-91254179).

## DISCUSSION

Modelling alternative sequence inclusion quantification from RNA-seq using the beta distribution allows the precision of its estimates to be proportional to the associated read coverage and reflected on the significance of differences in AS between samples. Besides the convenience of modelling precision, plotting the estimated beta distributions provides an intuitive graphical framework for understanding and evaluating the technical and biological uncertainties underlying PSI estimates. The main objective of differential AS analyses is to interpret the evidence, in terms of effect size and significance, collected for the difference in the inclusion of a given sequence of interest between biological conditions. As such, the `betAS` package was designed as a visual, flexible and easy-to-use decision-support tool to assist biologists analyse and understand their AS data.

As herein illustrated, a compromise between modelling the estimation uncertainty in the inclusion level quantification of individual samples and accounting for the variability among replicates is crucial for differential AS analysis, particularly when the sample size is small. Although several other methods using ΔPSI as a metric of the effect size have been proposed to address differential AS between conditions (18, 20), to our knowledge none has used the actual junction read counts to guide the user in visually interpreting PSI estimates and their confidence, even though these reads are proportional to the relative abundance of the alternative RNA sequences present in the profiled biological sample. `betAS` provides the first model that directly relies on the number of reads supporting inclusion or exclusion of alternative sequences to provide its users with an intelligible graphical assessment of the sources of uncertainty in AS analyses. `betAS` facilitates the visual interpretation of the effect size and significance statistics provided by differential AS tools, currently supporting the output processed files of `vast-tools` (16), `rMATS` (15), and `Whippet` (17), as they include inclusion and skipping read counts.

`betAS`‘ visual approach can be particularly helpful in interpreting the uncertainty in PSI estimates of AS events supported by very few reads. These events’ beta distributions are more dispersed than those for higher read counts and the variability in the PSI estimation is passed on to the comparison between groups, reflected on the respective significance quantification, and can be visually inspected via the shape of the beta distributions. `betAS` thereby enables users to visually assess if the samples’ read depth is enough for profiling the AS events of interest. This is particularly relevant for RNA-seq datasets with a small sample size, a large proportion of those generated and used to study specific splicing regulatory mechanisms, in which the impact of individual samples’ read coverage on the precision of differential AS analysis is stronger. Importantly, the implementation of the `betAS` pipeline as a web app allows any user, irrespectively of their familiarity with R or the command line, to perform differential AS analyses across two or more conditions.

While `betAS` harbours useful features for AS analysis, it also has inherent limitations. Firstly, `betAS` relies on read count data obtained from different AS analysis tools, each dependant on a mapping approach, whose accuracy impacts the precision of inclusion/exclusion calling on which PSI quantification depends. Importantly, this dependence on prior read count data implies that users are required to have minimum computational proficiency, to run sequence alignments. We compared, for the same sample, read count estimates by `vast-tools`, `rMATS` and `Whippet`, finding them overall generally (but not perfectly) consistent across tools (Supplementary Figure 11A-C). PSI quantifications are also generally consistent but we found extremely discrepant events (e.g., PSI of 0% from one tool and of 100% from another), mostly associated with very low coverage (Supplementary Figure 11D). Users are therefore advised to critically determine the mapping procedure that best matches their specific requirements, becoming familiar with the underlying assumptions guiding that choice, namely those associated with the aligner and the transcriptome annotation, to name a few.

One of the most challenging aspects of differential AS analysis is the assessment of the biological and phenotypical consequences of a given alteration in PSI. PSI estimates for each biological sample always reflect a sampling of the whole set of RNA molecules therein and relative isoform expression changes have different impacts on the PSI depending on its initial value (Supplementary Figure 12). While the intuitive visualisation provided by `betAS` is certainly helpful, an integrated framework to easily check the known biological implications of differential AS from available databases will contribute to a clearer analysis of the biological consequences of AS alterations and a more complete usability experience for the user.

In summary, we propose betAS as an important contribution to the alternative splicing research field, since it provides sound differential splicing analysis, even of RNA-seq datasets with low read coverage and/or small sample size, while guiding researchers to biologically relevant phenomena across multiple conditions (including time series) with its easy, visual and intelligible way of interpreting statistics supporting differential AS that are not commonly intuitively explored.

## DATA AVAILABILITY

`betAS` web app is publicly available at https://compbio.imm.medicina.ulisboa.pt/app/betAS. Its source code is available at https://github.com/DiseaseTranscriptomicsLab/betAS.

## ACKNOWLEDGEMENTS

We thank colleagues Manuel Irimia, Benilton Carvalho, Ramiro Magno, and all the members of the Disease Transcriptomics Lab at iMM for valuable discussions and suggestions on the manuscript. We also thank the very knowledgeable anonymous reviewers designated by *RNA* for their constructive and insightful suggestions and criticisms to the first version of this manuscript; their feedback greatly contributed to improved versions of both the article and `betAS` itself.

---

**Supplementary Figure 1.**
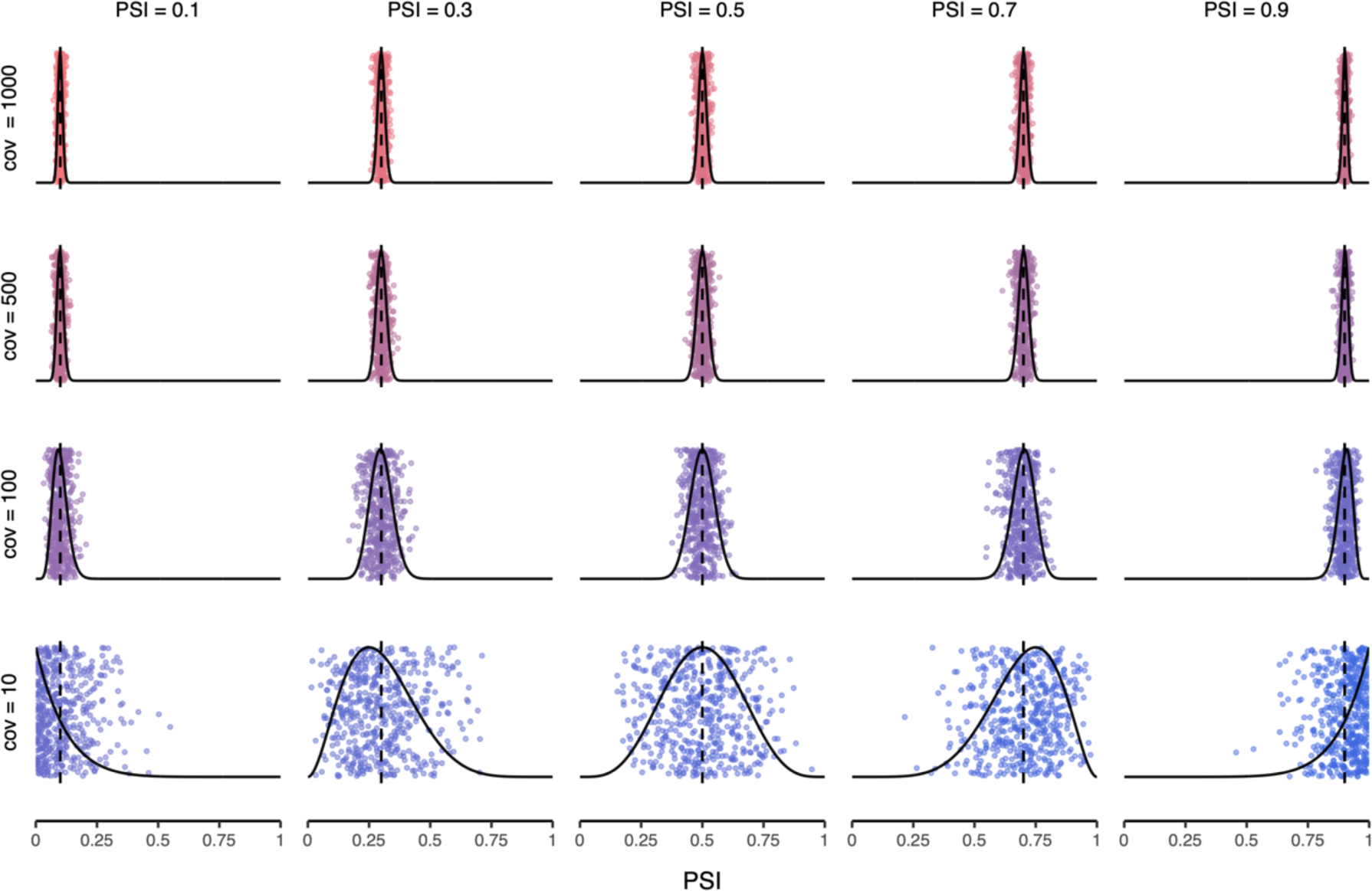
Beta distributions model PSI levels and associated confidence. Density plots (black lines) illustrating the dispersion of 500 values randomly generated from beta distributions (coloured vertically “jittered” points) modelling different PSI levels (columns) supported by different levels of coverage (cov) in junction read counts (rows).

**Supplementary Figure 2.**
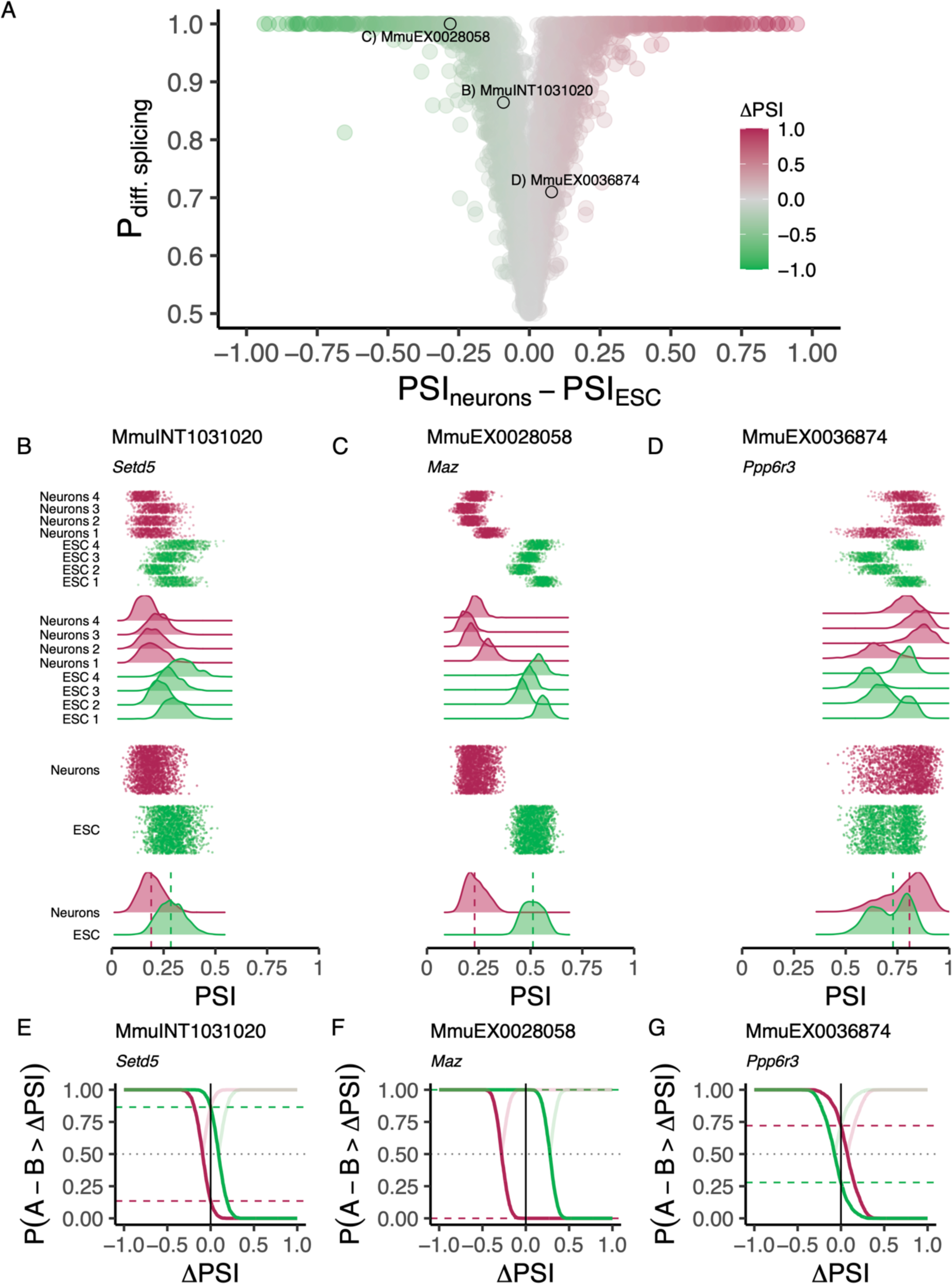
Estimating the probability of differential alternative splicing. (**A**) Volcano plot illustrating the effect size (ΔPSI) as the difference between the median group PSIs and their significance assessed by the estimated probability of differential AS, based on the proportion of differences between ESC and Neurons beta distribution randomly emitted values that are > 0 (P_diff_). Events are coloured by the ΔPSI calculated between neuronal and ESC samples. Marked events are illustrative of different scenarios of estimated P_diff_. (**B** to **D**) Beta distributions (vertical “jitter” and density plots of the emitted values) per sample (top) and per phenotypic group (bottom) and (**E** to **G**) cumulative line plots of the probability that the differences between values in sample groups A and B under comparison (A = ESC and B = neurons for the green line, with P(A – B > ΔPSI) reflecting P (ESC – neurons > ΔPSI); same for the red line, with A = neurons and B = ESC, to illustrate the symmetry in P_diff_ calculation) are greater than a given value of ΔPSI for selected events illustrative of different scenarios of estimated P_diff_: (**B,E**) MmuINT1031020 (gene *Setd5,* chr6:113109991-113110373), (**C,F**) MmuEX0028058 (gene *Maz*, chr7:127023482-127023706), (**D,G**) MmuEX0036874 (gene *Ppp6r3*, chr19:3544108-3544255). ESC: embryonic stem cells.

**Supplementary Figure 3.**
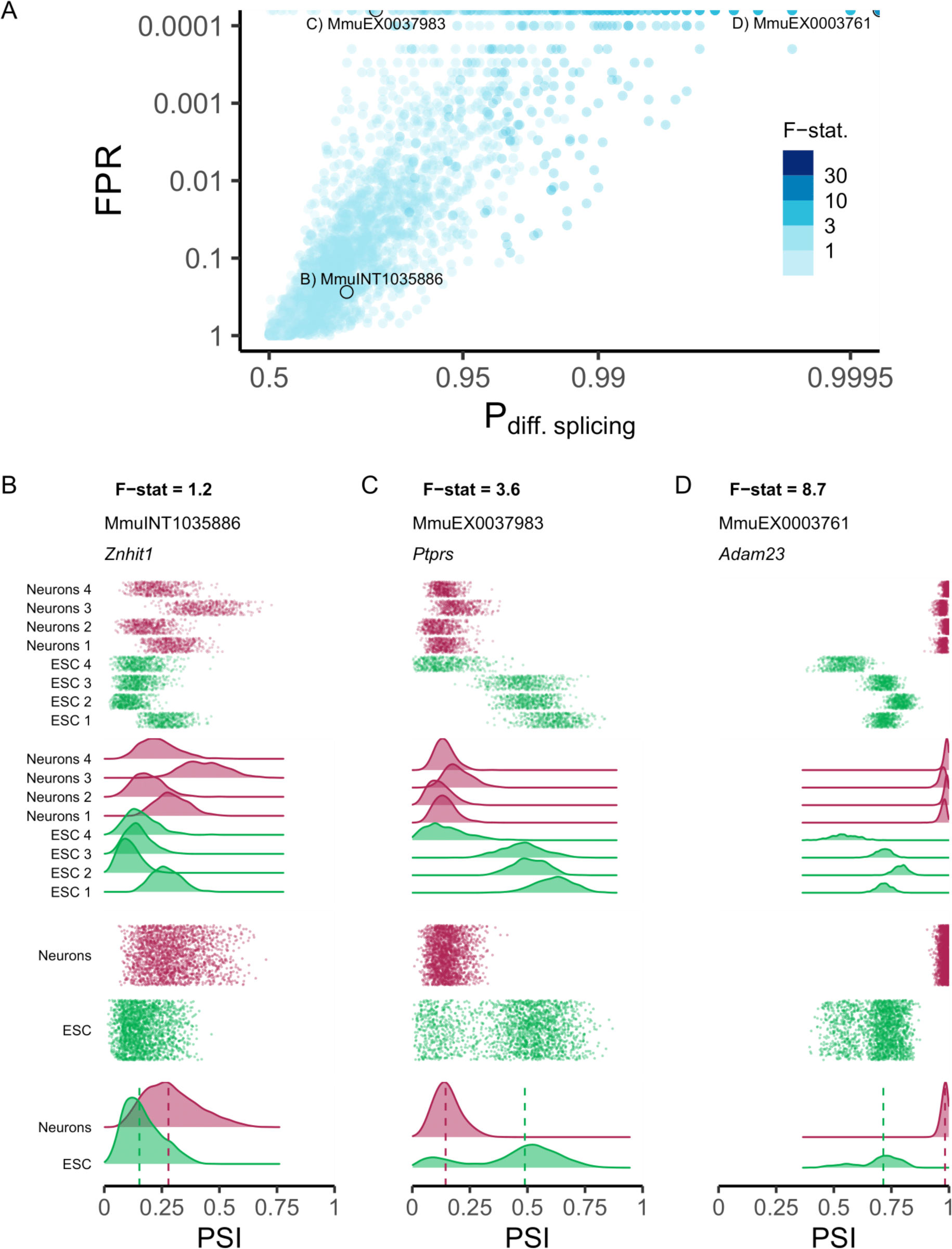
Estimating an F-statistic for differential alternative splicing. (**A**) Scatterplot comparing `betAS` metrics to estimate the significance of AS differences: false positive rate (FPR) and the probability of differential splicing, P_diff splicing_. Events are coloured by the F-statistic, i.e., the ratio of between- to within-group PSI variations. Marked events are illustrative of different scenarios of estimated P_diff_, FPR and F-statistic. Y-axis scales are log-transformed to facilitate visualisation. (**B** to **D**) Beta distributions (vertical jitter and density plots of the emitted values) per sample (top) and per group (bottom) of selected events illustrative of different scenarios of estimated F-statistic: (**B**) MmuINT1035886 (gene *Znhit1,* chr5:136985050-136987488), (**C**) MmuEX0037983 (gene *Ptprs*, chr17:56428852-56429157), (**D**) MmuEX0003761 (gene *Adam23*, chr1:63585365-63585455). ESC: embryonic stem cells.

**Supplementary Figure 4.**
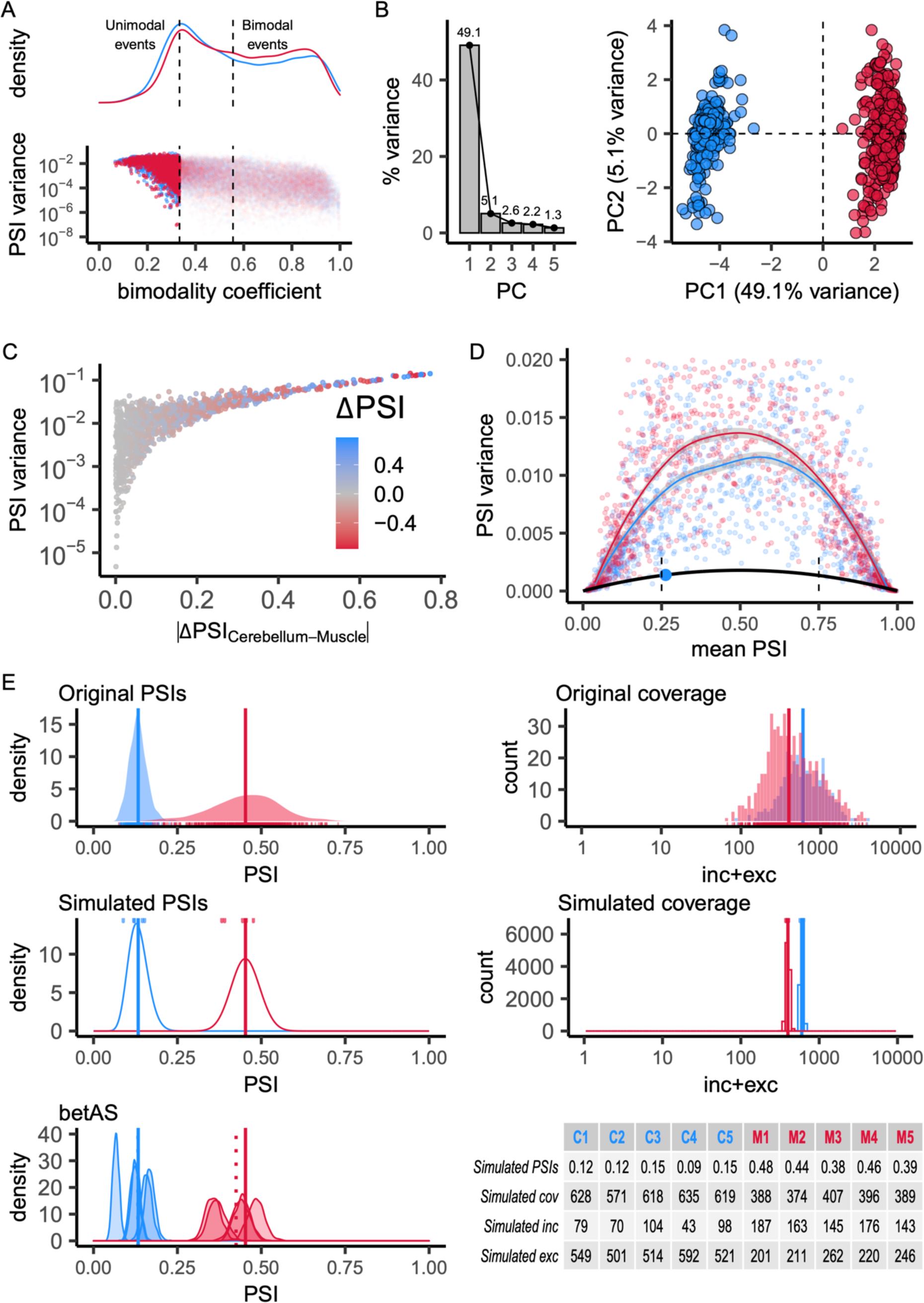
Simulation of empirically-inspired junction read counts. (**A**) Bottom: scatterplot of log-transformed PSI variance against the bimodality coefficient for considered events in muscle (red) and cerebellum (blue). Top: density plot of the bimodality coefficient for muscle and cerebellum, illustrative of unimodal and bimodal subpopulations. Vertical dashed lines indicate bimodality coefficients cut-offs: events were considered unimodal if bimodality coefficient < 3/9 and bimodal if bimodality coefficient > 5/9. (**B** to **D**) Subset of unimodal events only. (**B**) Principal component analysis (PCA) performed on the centred, unscaled original PSI values. Left: Percentage of variance explained by each of the five first principal components (PC). Right: Scatterplot of samples’ loadings on PCs 1 and 2. (**C**) Scatterplot of log-transformed PSI variance against the ΔPSI between cerebellum’s and muscle’s PSI_REF_. (**D**) Scatterplot of PSI variance to PSI mean relationship for a subset of low variance events (PSI variance < 0.02), allowing the empirical determination of the lowest PSI variance (the least biological noise) within considered PSI values (point highlighted in blue) and the respective PSI variance/mean relationship (black curve). Vertical dashed lines indicate the mean PSI range considered to identify a subset of representative low variance events. (**E**) Illustrative example of the procedure to simulate RNA-seq junction read counts based on simulated PSIs and coverage on which `betAS` is applied. “Original PSIs”: density and rug plot showing the original distribution of PSI values for all samples; vertical lines indicate PSI_REF_ values for muscle and cerebellum. “Original coverage”: histogram and rug plot showing the original distribution of coverage values for all samples; vertical lines indicate cov_REF_ values for muscle and cerebellum. “Simulated PSIs”: density plot showing the beta distribution from which low variance PSI values around the mean of PSI_REF_ (vertical lines) are simulated (top rug plot). “Simulated coverage”: histogram showing the Poisson distribution from which coverage values around cov_REF_ (vertical lines) are simulated (top rug plot). “betAS”: density plots representing PSI estimation as done in `betAS`, with values randomly emitted from a beta distribution with the shape associated with the simulated junction read counts per sample. Table compiling simulated PSI, coverage and junction read count (inc and exc) values for the 5 simulated cerebellum (C1 to C5) and muscle (M1 to M5) replicates. Solid vertical lines indicate PSI_REF_ values for muscle and cerebellum, while dashed vertical lines indicate the median PSI values estimated by `betAS` for muscle and cerebellum.

**Supplementary Figure 5.**
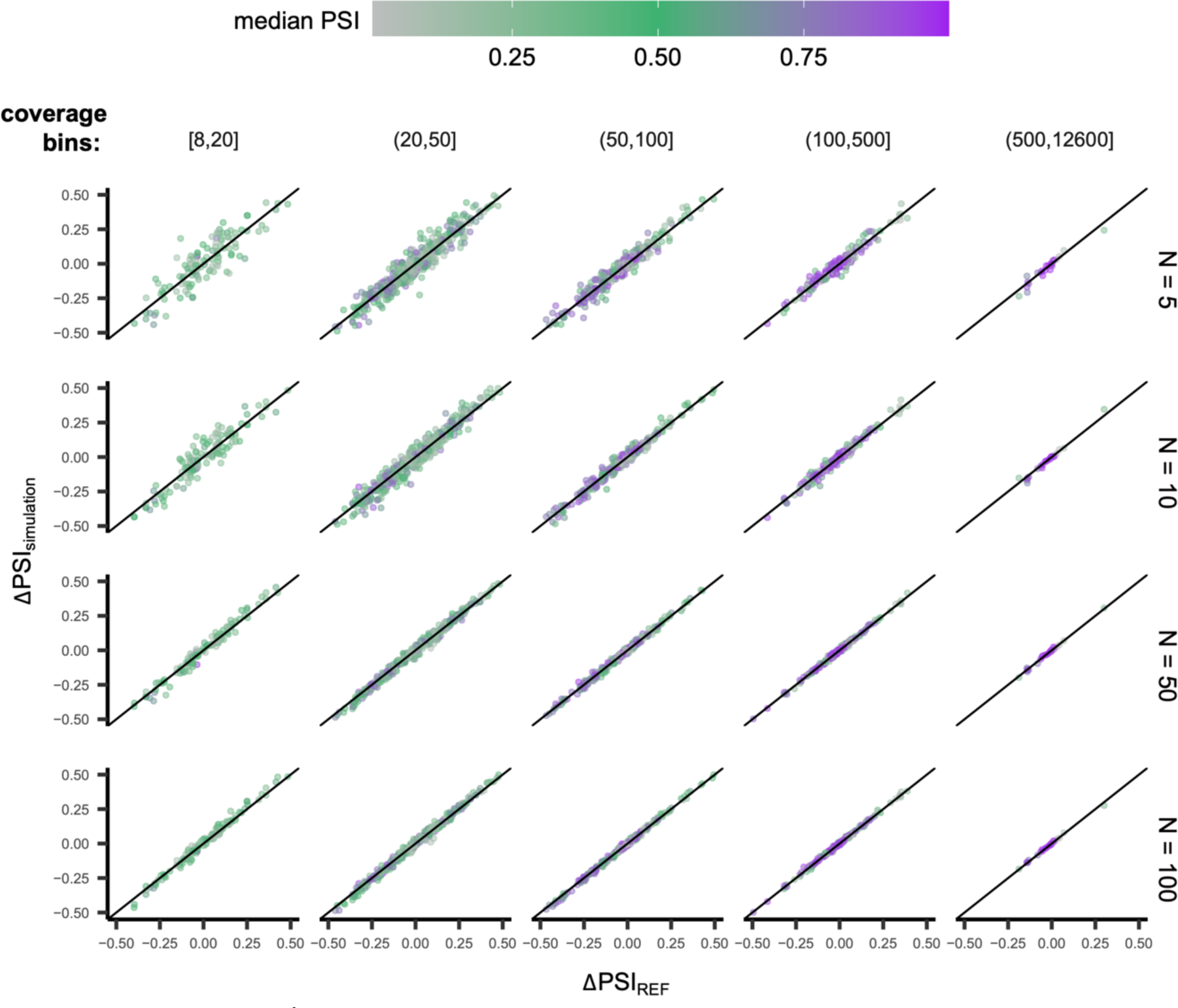
`betAS` accuracy in measuring PSI differences from simulated read counts. Scatterplots comparing the ΔPSI obtained by applying `betAS` to the simulated junction read counts, ΔPSI_simulation_, with the reference ΔPSI_REF_ for the subset of unimodal events considered, for different numbers of replicates (N, rows) and coverage ranges (in junction read counts, columns).

**Supplementary Figure 6.**
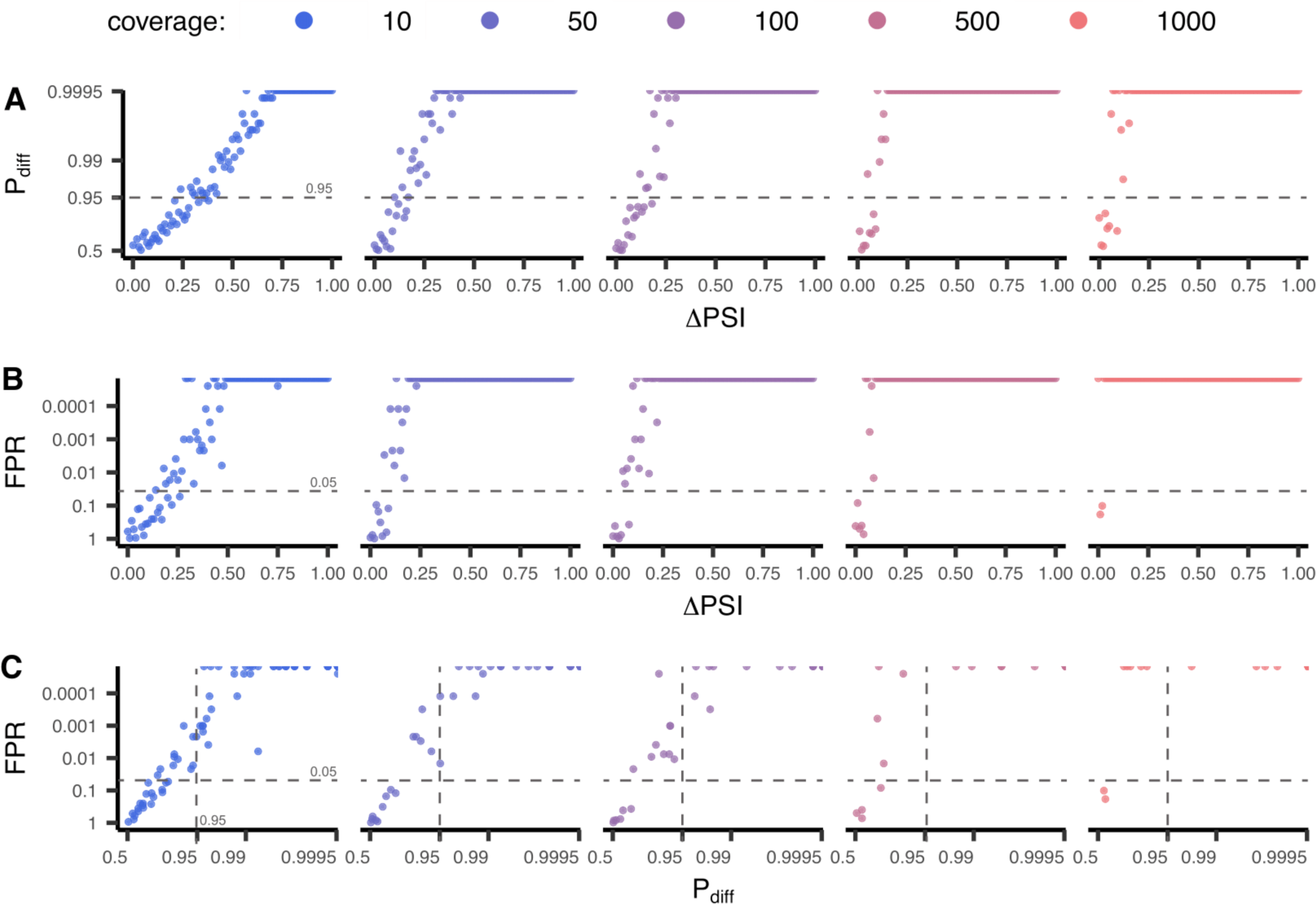
The impact of simulated cases of effect size (ΔPSI) and coverage pairs in P_diff_ (**A**) and FPR (**B**). P_diff_ and FPR values (directly compared in **C**) obtained for the differential splicing analysis comparing three biological replicate PSI values. Simulated PSI values were obtained considering ΔPSI ranging from 0 to 1 in intervals of 0.01 and coverage of 10, 50, 100, 500 or 1000 read counts. For each ΔPSI and coverage pair, two random PSI values were obtained with a difference of ΔPSI; for each of these PSI values, biological PSI triplicates were obtained by emitting 3 values from a beta distribution with parameters such that α + β = 138.5042 (see Supplementary Methods).

**Supplementary Figure 7.**
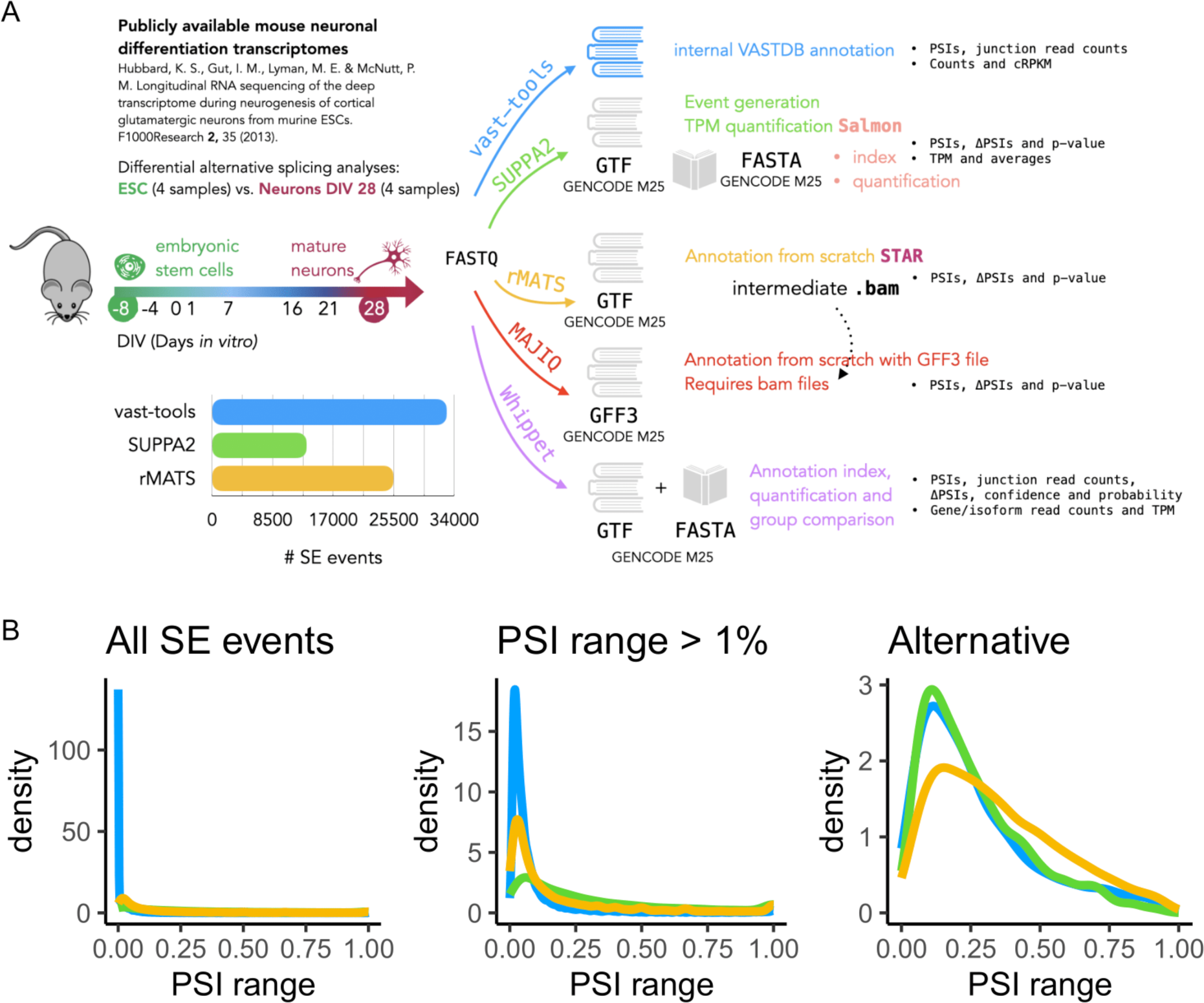
Comparison of differential alternative splicing analysis tools. (**A**) Explanatory diagram of the approach to obtain read counts and differential AS quantification metrics for each studied tool. (**B**) Density plots of the distribution of PSI for all exon skipping (SE) events considered (left), those with a PSI range (maximum to minimum PSI) > 1% (middle) and alternative events (0 < PSI < 1, right) for `vast-tools` (blue), `SUPPA` (green) and `rMATS` (yellow).

**Supplementary Figure 8.**
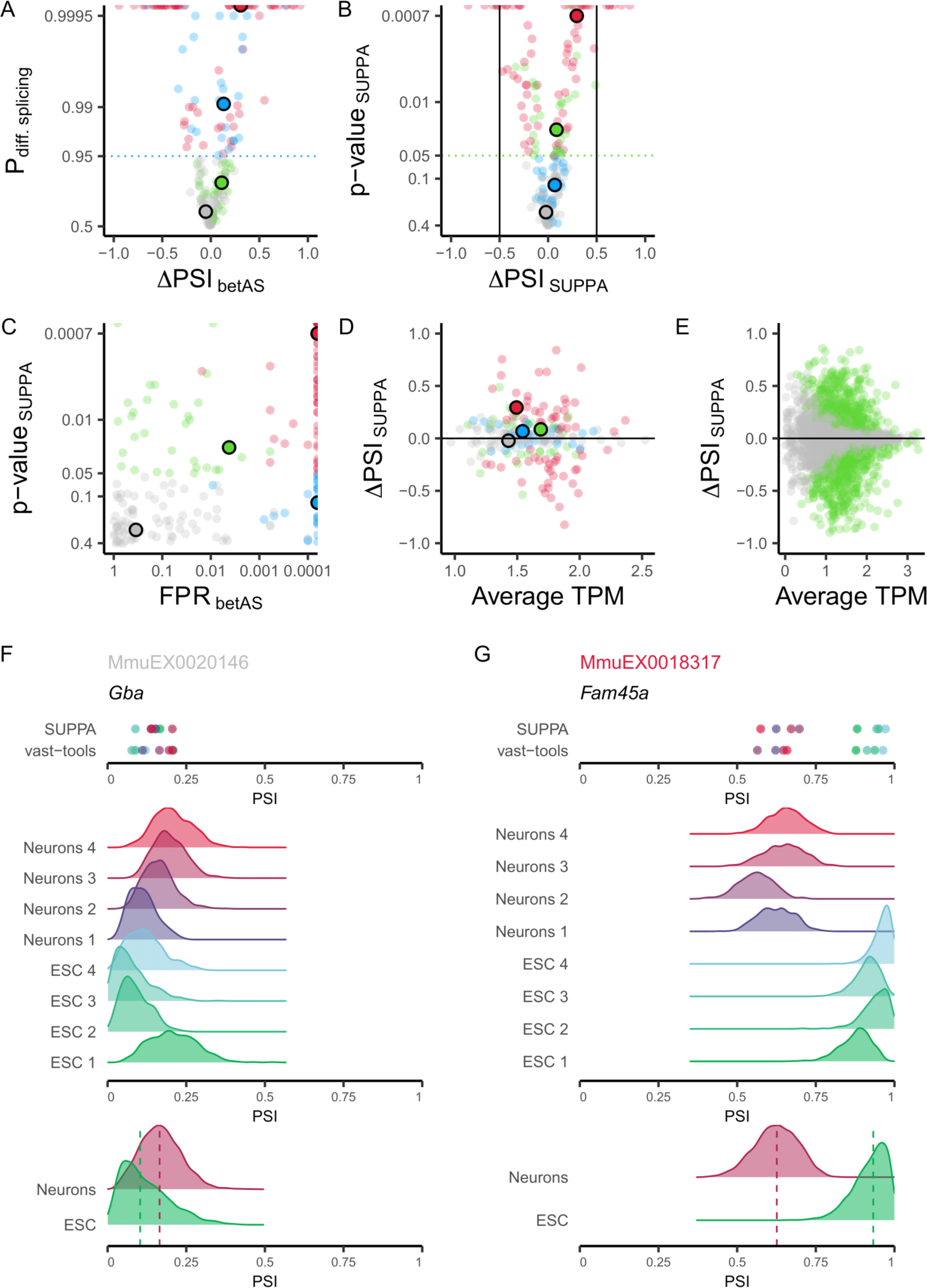
Comparison of differential AS between `betAS` and `SUPPA`. (**A**) `betAS`’ volcano plot of differential AS (each AS event represented by a dot) illustrating the effect sizes (ΔPSI_betAS_) as the differences between the median group PSIs and their significance assessed by the estimated probability of differential AS, based on the proportion of differences between beta distributed randomly emitted values per sample group that are > 0 (P_diff. splicing_). The blue dashed horizontal line indicates the considered cut-off of P_diff_ = 0.95. (**B**) `SUPPA`’s volcano plot of differential AS illustrating the effect sizes (ΔPSI_SUPPA_) and their significance assessed by `SUPPA`’s p-value. The green dashed horizontal line indicates the considered cut-off of p-value = 0.05. (**C**) Scatterplot comparing `betAS`’ and `SUPPA`’s significance metrics for differential AS: `betAS`’ false positive rate (FPR) and `SUPPA`’s p-value. Points are coloured based on the differential AS calls by both tools as in Figure 4: events considered differentially spliced by both `betAS` and `SUPPA` (red), by `betAS` (blue) or `SUPPA` (green) alone or by none of the tools (grey). (**D** and **E**) Scatterplots comparing ΔPSI_SUPPA_ with the respective transcript’s average expression in transcripts per million (TPM) for the subset of events analysed between `betAS` and `SUPPA` (**D**) and for all `SUPPA` events (**E**). (**E**) events highlighted in green are those called differentially spliced by `SUPPA` (p-value < 0.05). (**F** and **G**) `SUPPA` and `vast- tools`’ PSIs (top) and densities of emitted beta distributed values for individual samples (middle) and their merging per sample group (bottom, with dashed lines signing median values) for selected events illustrative of different combinations of effect size and significance of AS differences, identified by `vast-tools`’ IDs (VAST-DB annotation for the mouse mm10 genome assembly): (**F**) MmuEX0020146 (gene *Gba,* chr3:89203192-89203360), (**G**) MmuEX0018317 (gene *Fam45a*, chr19:60817531-60817610). ESC: embryonic stem cells.

**Supplementary Figure 9.**
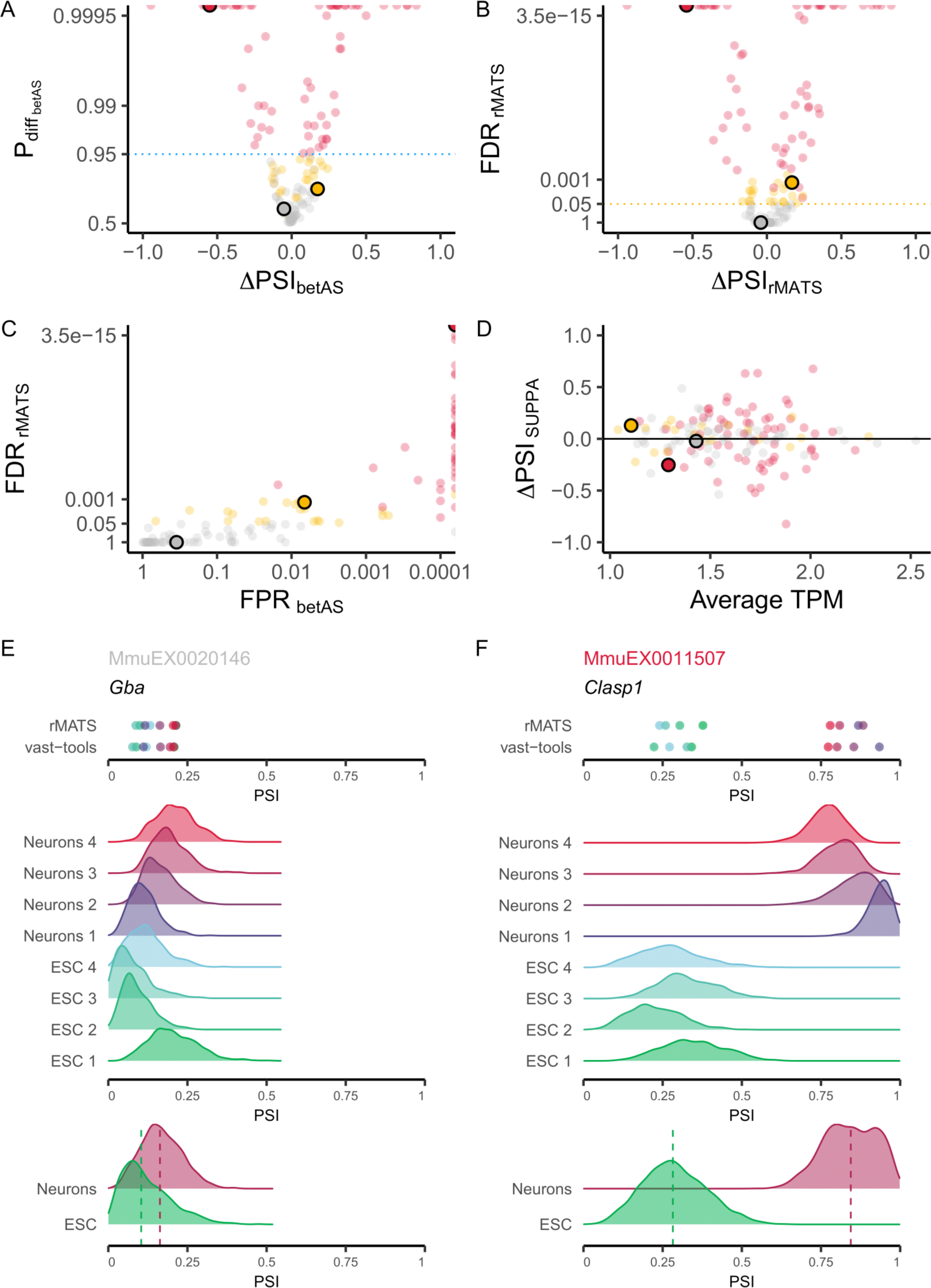
Comparison of differential AS between `betAS` and `rMATS`. (**A**) `betAS`’ volcano plot of differential AS (each AS event represented by a dot) illustrating the effect sizes (ΔPSI_betAS_) as the differences between the median group PSIs and their significance assessed by the estimated probability of differential AS, based on the proportion of differences between beta distributed randomly emitted values per group that are > 0 (P_diff betAS_). The blue dashed horizontal line indicates the considered cut-off of P_diff_ = 0.95. (**B**) `rMATS`’s volcano plot of differential AS illustrating the effect sizes (ΔPSI_rMATS_) and their significance assessed by `rMATS`’s FDR. The yellow dashed horizontal line indicates the considered cut-off of FDR = 0.05. (**C**) Scatterplot comparing `betAS`’ and `rMATS`’s significance metrics for differential AS: `betAS`’ false positive rate (FPR) and `rMATS`’ FDR. Points are coloured based on the differential AS calls by both tools as in Figure 4: events considered differentially spliced by both `betAS` and `rMATS` (red), by `betAS` (blue) or `rMATS` (yellow) alone or by none of the tools (grey). (**D**) Scatterplots comparing ΔPSI_SUPPA_ with the respective transcript’s average expression in transcripts per million (TPM) for the subset of events analysed between `betAS`, `rMATS` and `SUPPA`. (**E** and **F**) `rMATS` and `vast-tools`’ PSIs (top) and densities of emitted beta distributed values for individual samples (middle) and their merging per sample group (bottom, with dashed lines signing median values) for selected events illustrative of different combinations of effect size and significance of AS differences, identified by `vast-tools`’ IDs (VAST-DB annotation for the mouse mm10 genome assembly): (**E**) MmuEX0020146 (gene *Gba,* chr3:89203192-89203360), (**F**) MmuEX0011507 (gene *Clasp1*, chr1:118541675-118541698). ESC: embryonic stem cells.

**Supplementary Figure 10.**
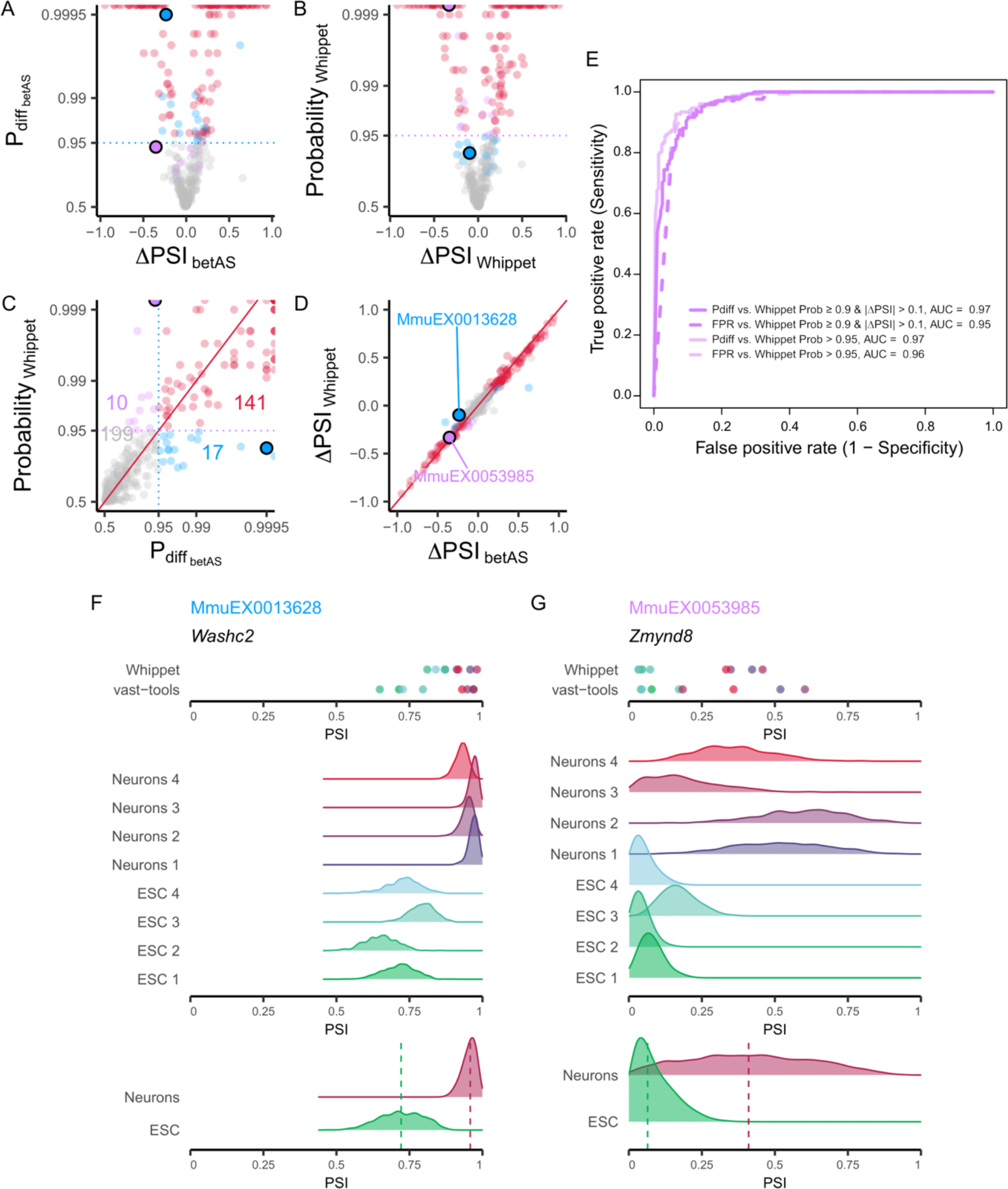
Comparison of differential AS between `betAS` and `Whippet`. (**A**) `betAS`’ volcano plot of differential AS (each AS event represented by a dot) illustrating the effect sizes (ΔPSI_betAS_) as the differences between the median group PSIs and their significance assessed by the estimated probability of differential AS, based on the proportion of differences between beta distributed randomly emitted values per sample group that are > 0 (P_diff. splicing_). The blue dashed horizontal line indicates the considered cut-off of P_diff_ = 0.95. Points are coloured based on the differential AS calls by both tools: events considered differentially spliced by both `betAS` and `Whippet` (red), by `betAS` (blue) or `Whippet` (purple) alone or by none of the tools (grey). Selected example events are depicted as larger outlined dots. (**B**) `Whippet`’s volcano plot of differential AS illustrating the effect sizes (ΔPSI_Whippet_) and their significance assessed by `Whippet`’s “probability” (probability that there is some change in ΔPSI, given the read depth of the AS event). The purple dashed horizontal line indicates the considered cut-off of probability = 0.95. Points are coloured based on the differential AS calls by both tools and selected example events are depicted as larger outlined dots. (**C**) Scatterplot comparing `betAS`’ and `Whippet`’s differential AS significance metrics. `betAS`’ estimated probability of differential AS, based on the proportion of differences between the beta distributed randomly emitted values per group that are > 0 (P_diff_) and `Whippet`’s probability, with dashed lines defining quadrants indicating the significance cut-offs considered (P_diff_ > 0.95 for `betAS`, probability > 0.95 for `Whippet`). Red diagonal solid line indicates identity: P_diff betAS_ = Probability_Whippet_. Points are coloured based on the differential AS calls by both tools and selected example events for each quadrant as larger outlined dots. (**D**) Scatterplot comparing `betAS`’ and `Whippet`’s differential AS effect size (ΔPSI). Red diagonal solid line indicates identity: ΔPSI_Whippet_ = ΔPSI_betAS_. (**E**) Receiving Operating Characteristic (ROC) curves `betAS`’ differential AS calls with P_diff_ (solid lines) and FPR (dashed lines), considering as ground truth the differential calls from `Whippet` (dark purple: Probability ≥ 0.9 and |ΔPSI| > 0.1; light purple: Probability > 0.95). (**F** and **G**) `Whippet` and `vast- tools`’ PSIs (top) and densities of emitted beta distributed values for individual samples (middle) and their merging per sample group (bottom, with dashed lines signing median values) for selected events illustrative of different combinations of effect size and significance of AS differences, identified by `vast-tools`’ IDs (VAST-DB annotation for the mouse mm10 genome assembly): (**F**) MmuEX0013628 (gene *Washc2,* chr6:116256227-116256289), (**G**) MmuEX0053985 (gene *Zmynd8*, chr2:165852868-165852879). ESC: embryonic stem cells.

**Supplementary Figure 11.**
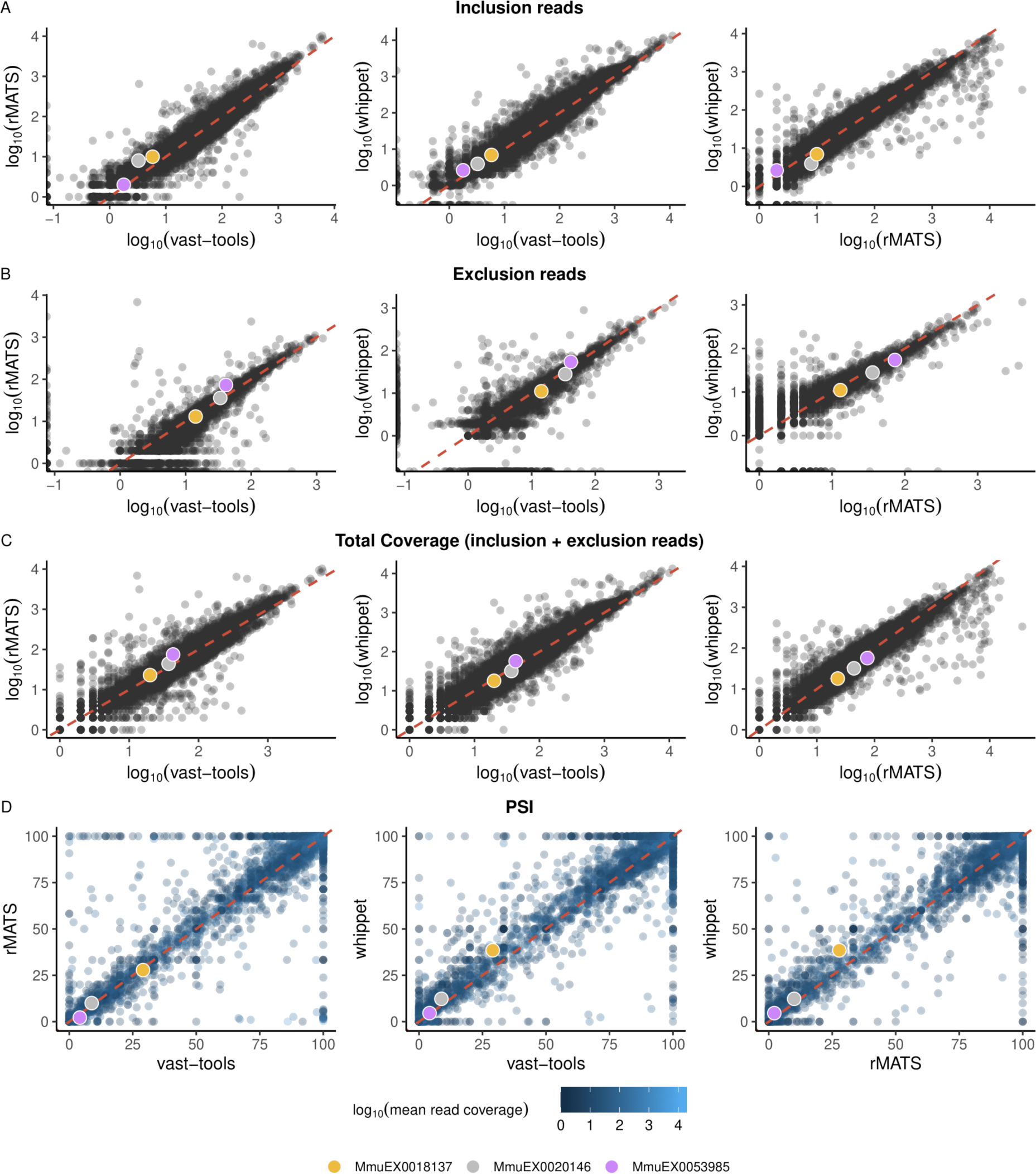
Impact of AS tools on event-supporting read counts and PSI values. Scatter plots depicting pairwise comparisons of alternative splicing tools supported by betAS (vast- tools, rMATS, and Whippet) for (**A**) inclusion reads, (**B**) exclusion reads, (**C**) total coverage (sum of inclusion and exclusion reads), and (**D**) Percent Spliced-In (PSI) values within the [0,100] interval. This analysis uses sample SRR645826 from the mouse neuronal differentiation timeline dataset as an example. PSI values (D) are color-coded based on the mean coverage between the tools being compared. Left plots compare vast-tools with rMATS, middle plots compare vast-tools with Whippet, and right plots compare rMATS with Whippet. Events displayed for each pairwise comparison are those shared between the two tools, identified by their genomic coordinates. Colored dots highlight three exemplary events discussed in the manuscript.

**Supplementary Figure 12.**
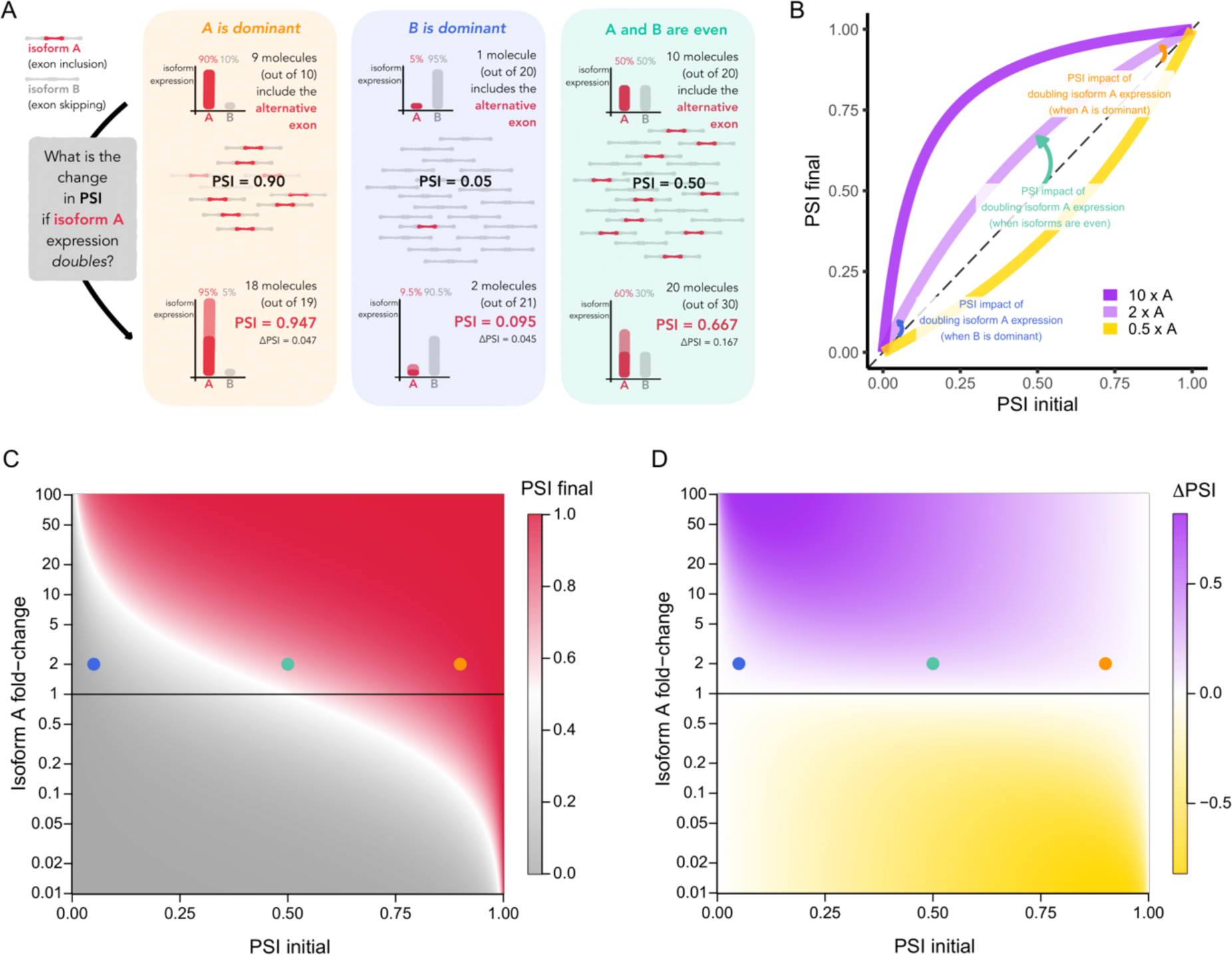
The impact of isoform expression changes on the PSI. (**A**) Explanatory diagram illustrative of the impact on the PSI of a duplication in the expression levels of the inclusion isoform in three different scenarios: when A is the dominant isoform (left), when the B isoform is the dominant (middle) and when A and B are evenly distributed. (**B**) Line plots illustrating PSI transitions (i.e., final PSI (Y-axis) against the initial PSI (X-axis)) associated with selected fold-changes of an inclusion isoform. The transitions shown in the three scenarios depicted in panel A are marked as arrows in the respective colours. (**C**) Heatmap showing the continuous spectrum of PSI transitions (i.e. final PSI (heat colour) against the initial PSI (X-axis)) depending on the A isoform fold-change (Y-axis). The transitions shown in the three scenarios depicted in panel A are marked with dots in the respective colours. (**D**) Heatmap showing the continuous spectrum of ΔPSI (final PSI – initial PSI, heat colour) values for PSI transitions depending on the initial PSI (X-axis) and A isoform fold-change (Y-axis). The transitions shown in the three scenarios depicted in panel A are marked with dots in the respective colours.

