## Supplementary Materials for "betAS: intuitive analysis and visualisation of differential alternative splicing using beta distributions"

### SUPPLEMENTARY METHODS

#### The beta and the binomial distributions

If we consider the set of inc + exc junction reads supporting a particular alternative splicing (AS) event, each single read can be interpreted as the result of a sampling experiment with two possible outcomes: either to support inclusion (considered as "success", with a given probability) or exclusion (a "failure"). A problem in which there is either success or failure, with a given probability of success, is a binomial problem. When the parameter of a binomial distribution (i.e., the probability of success) is not known, a beta distribution can be used as a general distribution of possible hypotheses for the probability value we want to estimate, as beta distributions are conjugate priors to the binomial likelihood (1). Thus, the *number of successes* inc can be modelled as a binomial distribution with the probability of success PSI as its parameter. Such PSI can be estimated using the beta distribution (Figure 1A), used to model proportions. The shape of the beta distribution is determined by two parameters, α and β, with its mean given by the ratio between α and α + β, which is directly comparable to PSI ratios, if α ~ inc and β ~ exc.

$$\text{mean of the beta distribution with parameters } \alpha \text{ and } \beta = \frac{\alpha}{\alpha + \beta}$$

For a given mean, the higher α and β are, the narrower the distribution is (Figure 1, right). Thus, shape parameters α and β can conveniently incorporate information on the number of reads used in the quantification.

#### Publicly available RNA-seq datasets used to illustrate the `betAS` differential AS analysis pipeline

We illustrate the `betAS` pipeline for the two-group comparison by analysing mouse embryonic stem cells (ESC) and differentiated cortical glutamatergic neurons (Neurons), making use of public RNA sequencing data (SRA project PRJNA185305) that profile transcriptomic changes during mouse neuronal differentiation (2). For the multiple group comparisons, we made use of the human subset of an RNA-seq time-series experiment on the transcriptomic basis of the development of seven major organs (ArrayExpress accession code E-MTAB-6814) (3).

#### `betAS` pipeline to model inclusion level quantification and differential AS

`betAS` requires an association between *samples* and *groups* (i.e., biological conditions), reflecting the experimental design. The pipeline starts by taking the inclusion (inc) and exclusion (exc) read counts associated with each AS event for each *sample* and uses function *rbeta* from the *stats* R package to emit a set of $N_{ind}$ values from a beta distribution with shape parameters α = inc and β = exc if both inc and exc are non-zero. `rbeta(n = N_ind, shape1 = α, shape2 = β)` exemplifies R code to generate **n** random values from a beta distribution with shape parameters **`shape1`** and **`shape2`**.

Given that, in most biological samples, the majority of annotated alternative sequences are actually either nearly always included (PSI ≈ 1) or nearly always excluded (PSI ≈ 0), empirical PSI distributions are typically zero-one-inflated:

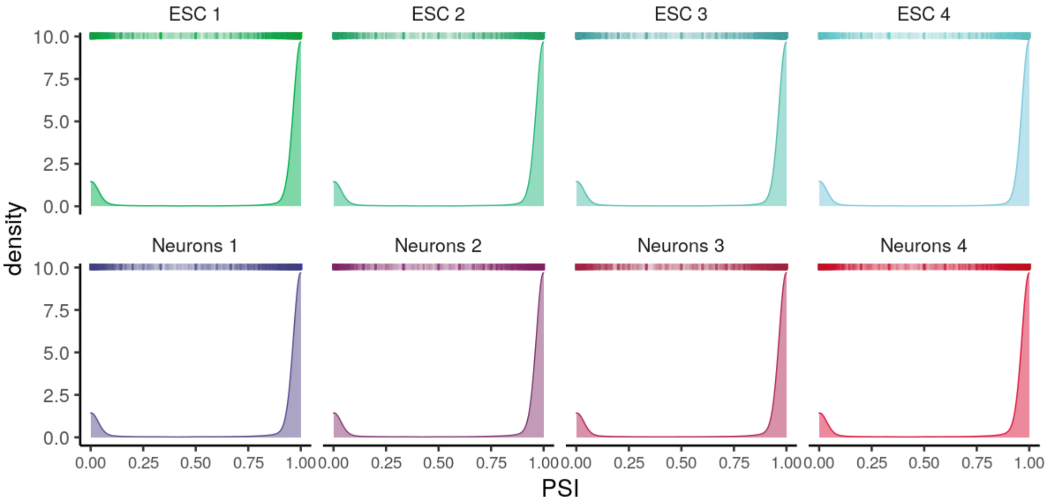

**Figure SM1: Empirical PSI distributions are zero-one-inflated:** most observed PSIs are close to 0 or 1. Density and rugplots (top) for the distribution of PSI values per sample for mouse embryonic stem cells (ESC) and differentiated cortical glutamatergic neurons (Neurons) that are part of the public RNA sequencing dataset PRJNA185305 (2).

While inc or exc can be equal to zero, values obtained with *rbeta* from a beta distribution with α = 0 or β = 0 are single-valued as 0 or 1, respectively, and are inappropriate to model inclusion levels. To account for this, different strategies were studied to implement an "increment" factor to be added to values of inc or exc equal to zero, so that the described beta distribution-based approach is still appropriate to model inclusion levels, i.e., such that the dispersion of the emitted values decreases as coverage increases but their average values do not deviate much from 0 or 1.

**incr = 1**

Since the minimum unit of evidence for a sequence is one read, the simplest way to increment inc or exc in order to avoid zeros would be to make α = 1 (when the measured PSI = 0) or β = 1 (when the measured PSI = 1). Incrementing in one the measured number of reads has an impact on the PSI deviation that is inversely proportional to the read coverage and observed to be too high when the coverage is of only a few reads (Figure SM2 below).

**incr = cov/99**

As an alternative, one could consider a coverage-dependent dynamic increment such that a PSI = 0 will be corrected to be 0.01 (an arbitrarily set value for the minimum biological PSI that is not zero):

$$\frac{\text{incr}}{\text{incr} + \text{cov}} = 0.01 \Leftrightarrow \text{incr} = \frac{\text{cov}}{99}$$

The increment would then be 1 for coverage ≥ 99. However, PSI estimates supported by very low coverage get an artificially high precision (Figure SM2 below), while PSI dispersion should decrease as coverage increases. We therefore tried instead a non-parametric approach based on the median of emitted values.

### Ad-hoc empirically determined incr

For a measured PSI of 0 or 1, we established the increment to inc or exc to be lower than one, as one read is the minimum unit of evidence for a sequence. Moreover, the increment should be such that the median of the beta distribution is arbitrarily limited to ≤ 0.01 or ≥ 0.99, respectively. For each coverage/increment pair, with coverage values being all integers between 1 and 100 reads and increments incr equally spaced by 0.01 between 0 and 1, 100 beta distributions of 500 values were generated, with parameters α = incr and β = coverage, to find, for each coverage, the maximum increment factor such that no more than 5 simulated distributions had a median higher than 0.01.

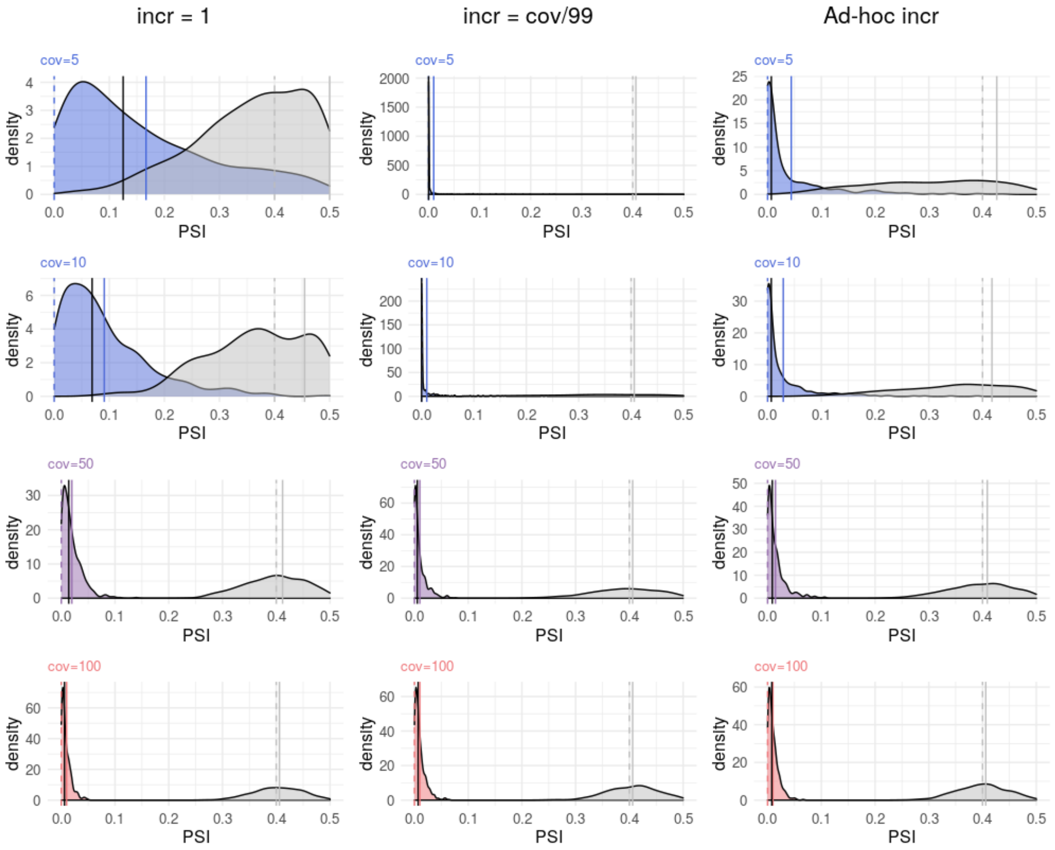

**Figure SM2: Comparison of the impacts on PSI between increment approaches**
In colour: beta distributions of PSI = 0 for each coverage in read counts (cov), after summing *incr* reads to the number of inclusion reads to avoid inc = 0; grey: beta distributions of PSI = 0.4 for each coverage, after summing *incr* to the number of inclusion reads; vertical dashed line: reference PSI (coloured for PSI = 0, grey for PSI = 0.4); vertical solid line: mean PSI after summing *incr* reads (coloured for PSI = 0, grey for PSI = 0.4); solid black line: median PSI after summing *incr* reads for PSI = 0. Left: impact of summing 1 to zero-valued inc read counts. Middle: impact of summing cov/99 reads to zero-valued inc read counts. Right: impact of summing an ad-hoc empirically determined incr (see description in this document's main text) to zero-valued inc read counts.

We empirically observed that this is approximately equivalent to considering a linear relationship between incr and coverage: incr = 0.01 · cov + 0.19 (Figure SM3). Moreover, we observe the expected decrease in PSI dispersion as coverage increases (Figure SM2).

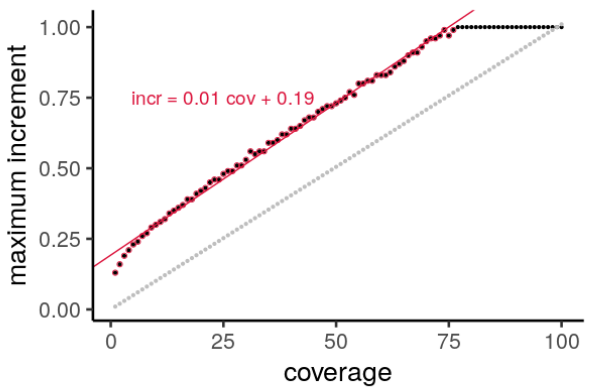

**Figure SM3: Empirical determination of the maximum increment as a function of coverage**
Scatterplot showing the maximum increment found per coverage (black) as well as the dynamic increment corresponding to the incr = cov/99 approach (grey). Linear regression equation and line for the empirically obtained values with a maximum increment below 1 (red).

After this procedure, for each event, sets of $N_{ind}$ *rbeta*-emitted values per sample are grouped per condition (ESC and Neurons in Figure 2) with the median per group as their summary average PSI (Figure 2C-D). The effect size of differential inclusion of the AS sequence between conditions is represented by ΔPSI:

$$PSI_{betAS}\text{ (condition)} = \textbf{median}\ (\texttt{rbeta}\ \mathbf{1}, \texttt{rbeta}\ \mathbf{2}, \ldots, \texttt{rbeta}\ \mathbf{N})$$

(where `rbeta` **i** represents the set of rbeta-emitted values for sample i)

$$\Delta PSI = PSI_{betAS}\ (\textbf{ESC}) - PSI_{betAS}\ (\textbf{Neurons})$$

Analogously to the `vast-tools` *diff* module (4), the `betAS` pipeline takes the proportion of differences between the two shuffled sets of emitted values per condition that are greater than a minimum value (minDiff) as the significance metric for each estimated ΔPSI (Supplementary Figure 2). Using minDiff = 0 has the same interpretation as asking what proportion of *rbeta*-emitted values for ESC are higher than those emitted for Neurons, thus reflecting the probability of differential splicing, $P_{x=0}$ of $PSI_{betAS}$ (**ESC**) being greater than $PSI_{betAS}$ (**Neurons**):

$$P_{x=0} = \frac{\#\ (\textbf{differences}\text{ that are} > 0)}{\#\text{ all }\textbf{differences}}$$

with **differences** = emitted values $_{\textbf{ESC}}$ - emitted values $_{\textbf{Neurons}}$

`betAS` allows users to tune the significance of estimated ΔPSIs, by setting the minimum difference minDiff in order to calculate $P_{x=minDiff}$:

$$P_{x=minDiff} = \frac{\#\ (\textbf{differences}\text{ that are} > \text{minDiff})}{\#\text{ all }\textbf{differences}}$$

with **differences** = emitted values $_{\textbf{A}}$ - emitted values $_{\textbf{B}}$

This calculation is performed spanning a set of ΔPSI values between -1 and 1 with increments of 0.002 (Supplementary Figure 2E-G). In a comparison between two groups with different number of samples, the set of *rbeta*-emitted values from the largest group is sampled to match the dimension of the vector representing the smaller group.

Importantly, since the comparison is sided (in this case, emitted values $_{\textbf{ESC}}$ - emitted values $_{\textbf{Neurons}}$ > 0), both low and high probabilities reflect the most extreme differences. For instance, with minDiff = 0 a $P_{x=0}$ = 0.8 would reflect that there is a 0.8 probability of higher inclusion values in ESC than in Neurons (while the probability of Neurons having greater inclusion values than ESC is 0.2) but a probability of $P_{x=0}$ = 0.2 would reflect the opposite case, with the probability of inclusion levels in ESC being greater than in Neurons equal to 0.2 and that of levels in Neurons being greater than in ESC equal to 0.8. Accordingly, to simplify the interpretation of this metric, betAS transforms the values of $P_{x=0}$ from the interval [0,1] to the interval [0.5,1]:

$$P_{diff} = |P_{x=0} - 0.5| + 0.5$$

This defines an intelligible significance metric that, coupled with the ΔPSI, allows an informed selection of interesting (according to the user's criteria) differentially spliced sequences between two conditions (Figure 2C-D).

**Pipeline to simulate empirically-inspired RNA-seq junction read counts**

With the purpose of simulating RNA-seq junction read counts, inspired in real PSIs and coverages, on which to test the accuracy of betAS in quantifying AS differences, we started by identifying a set of GTEx (5) studied cerebellum- and muscle-specific AS events reflecting the lowest biological variability, i.e., the least observed biological noise.
By studying the relationship between the PSI variance and mean for the subset of 1143 tissue-unimodal events (Supplementary Figure 4B-D) while considering the mean and variance relationship for a beta distribution with shape parameters α and β (equations **1** and **2**), a smaller set of representative events showing the lowest observed variance with PSI mean values ranging between 0.25 and 0.75 was considered to determine the combination of α and β that is equivalent to the lowest biological variance.

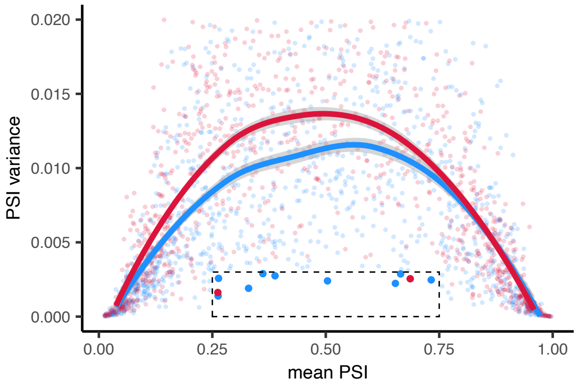

**Figure SM4: PSI variance and mean relationship for a subset of tissue-specific unimodal events**.
PSI variance plotted against PSI mean per event across muscle (red) or cerebellum (blue) GTEx samples. Curves show the local polynomial regression fitting ("loess") for all the 1143 events considered in muscle and cerebellum. The subset of events selected to feed the empirical estimate of the lowest observed variance are highlighted. Dashed rectangle indicates the region in which events were considered as representative of the lowest biological variance.

$$\textbf{PSI mean} = \frac{\alpha}{\alpha + \beta} \text{ (equation } \mathbf{1}\text{) and } \textbf{PSI variance} = \frac{\alpha \cdot \beta}{(\alpha + \beta)^2 \cdot (\alpha + \beta + 1)} \text{ (equation } \mathbf{2}\text{)}$$

From equations 1 and 2, it can be concluded that, for a fixed PSI value, its variance is inversely proportional to α + β:

$$\text{PSI variance} = \frac{\text{PSI mean} \cdot (1 - \text{PSI mean})}{\alpha + \beta + 1}$$

We therefore solved the above equations in order to find α and β for each of the set of low-variance representative events:

$$\alpha = \frac{\text{PSI mean}^2}{\text{PSI variance}} \cdot (1 - \text{PSI mean} - \text{PSI variance})$$

$$\beta = \frac{1 - \text{PSI mean}}{\text{PSI mean}} \cdot \alpha$$

Then, for these points, the maximum α + β, i.e., the minimum variance was determined.

| Mean | Variance | $\alpha$ | $\beta$ | $\alpha + \beta$ |
|---|---|---|---|---|
| 0.504 | 0.00240411 | 51.878 | 51.105 | 102.983 |
| 0.264 | 0.00257553 | 19.614 | 54.770 | 74.385 |
| 0.388 | 0.00274568 | 33.184 | 52.311 | 85.495 |
| 0.665 | 0.00287193 | 50.919 | 25.665 | 76.584 |
| 0.653 | 0.00223484 | 65.559 | 34.782 | 100.341 |
| 0.361 | 0.00289198 | 28.483 | 50.323 | 78.806 |
| 0.732 | 0.00247098 | 57.368 | 20.977 | 78.345 |
| 0.330 | 0.00190498 | 37.955 | 77.092 | 115.047 |
| **0.263** | **0.00138866** | **36.395** | **102.109** | **138.504** |

| Mean | Variance | $\alpha$ | $\beta$ | $\alpha + \beta$ |
|---|---|---|---|---|
| 0.262 | 0.00162246 | 30.993 | 87.229 | 118.222 |
| 0.686 | 0.00254829 | 57.299 | 26.217 | 83.517 |

Their PSI variance to PSI mean relationship matched that of a beta distribution with shape parameters such that α + β = 138.5042.

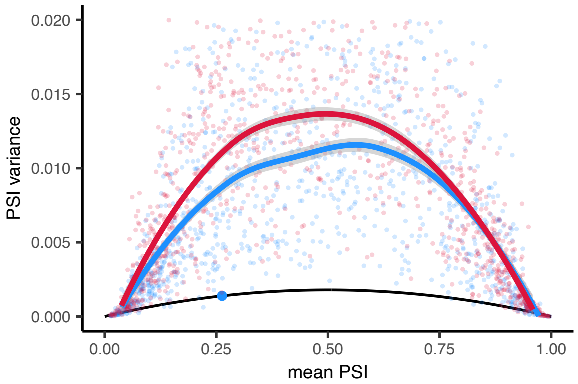

**Figure SM5: PSI variance and mean relationship for the low-variance events**. PSI variance plotted against PSI mean per event across muscle (red) or cerebellum (blue) GTEx samples. Curves show the local polynomial regression fitting ("loess") for all the 1 143 events considered in muscle and cerebellum. The relationship between PSI variance and mean that is associated with the lowest variance, with α + β = 138.5042, is marked by a black line, with the AS event associated with this maximum highlighted by the larger blue dot.

Then, for each event, beta distributions with the lowest empirical variance, i.e., such that $\alpha + \beta = 138.5042$, and with mean values equal to the reference PSI for that tissue/event pair ($PSI_{REF}$) were used to simulate N (intended number of simulated biological replicates) PSI values.

In R:

```
simulatedPSIs <- rbeta(n = N, shape1 = a, shape2 = b),
```

with $a + b = 138.5042$ and $\frac{a}{a+b} = PSI_{REF}$

The Poisson distribution is used to model the probability of a given number of events occurring in a given interval and serves as a standard model for count data under the assumptions that: 1) transcripts are generated in a sample at a given transcription rate; 2) transcript fragments are detected with a given probability; 3) the number of observed reads follows a Poisson distribution with a rate (parameter) that is proportional to the product between that transcription rate and the probability of fragment detection. Along the lines of the PSI value simulation described above, N coverage values were thus simulated from a Poisson distribution with parameter λ equal to the reference coverage of that tissue/event pair ($cov_{REF}$):

```
simulatedCoverages <- rpois(n = N, lambda = covREF)
```

The binomial distribution with parameters n and p is appropriate to model read counts because it reflects the number of events in a series of n independent Bernoulli experiments, each with the same probability of occurrence p (see section *The beta and the binomial distributions*). Then, by combining simulated PSIs and coverages, RNA-seq splice junction read counts are simulated from a binomial distribution using the *rbinom* function:

```
bin <- rbinom(n = cov, size = 1, prob = psi)
inc <- sum(bin)
exc <- cov - inc
```

**Visual comparison of approaches for differential splicing analyses between two conditions**

We compared the `betAS` approach with other tools for differential splicing analysis by making use of public RNA sequencing data (SRA project PRJNA185305) on mouse embryonic stem cells and differentiated cortical glutamatergic neurons. In order to illustrate, discuss and interpret the particularities of measuring the inclusion of alternative sequences using evidence from RNA-seq data, we considered tools that fulfil all the following criteria:

1. quantify differential AS between samples using FASTQ or BAM files;
2. provide a significance metric to assist the interpretation of splicing differences;
3. use PSIs (or equivalent) to quantify the inclusion of AS events.

With these criteria, we initially selected `SUPPA2` (6), `rMATS` (7), `Whippet` (8) and `MAJIQ` (9) to benchmark differential AS statistics provided by `betAS` applied over `vast-tools`' AS quantifications

(Supplementary Figure 7). Overall quality of raw RNA-seq data of ESC and mature 28-DIV neurons samples was assessed on the respective FASTQ files using `fastqc` version 0.11.5 (https://www.bioinformatics.babraham.ac.uk/projects/fastqc) with no relevant quality problem or technical bias identified.

`vast-tools` (10–12) version 2.5.1 was used for quantification of inclusion levels of the alternative sequences in VAST-DB annotation for the mouse mm10 genome assembly. To ensure sufficient and reliable read coverage, only events with PSIs quantified for all samples (i.e., with no NA values) and a minimum mappability-corrected read coverage score of "VLOW" across all samples were considered. Only exon skipping events were considered for simplicity and easier interpretability. Quality columns from `vast-tools` output table containing inclusion levels (".Q") were used to extract inc and exc mappability-corrected junction read counts to serve as input for `betAS`, for each quantified PSI per event and sample. `betAS` analysis based on `vast-tools` quantifications was performed as detailed in the previous sections.

`SUPPA2` was run as detailed in their GitHub tutorial (https://github.com/comprna/SUPPA/wiki/SUPPA2-tutorial). First, transcript quantification was performed using `salmon` version 1.3.0 indexed on mouse Gencode transcript annotation GENCODE M25 (gencode.vM25.transcripts.fa), followed by quantification in the quasi-mapping-based mode with library type automatically inferred (`-l A`) and `--gcBias` flag on. `SUPPA2` in-house "multipleFieldSelection" script was used (with index row `k=1` and column to extract `f=4`) to extract TPMs from salmon quant.sf files. Moreover, as explained in the tutorial, transcript IDs in the previous set's output file iso_tpm.txt were formatted to match those in the GTF file that `SUPPA2` uses to calculate inclusion levels.

Next, event generation was performed using GENCODE M25 annotation and `--pool-genes` option for all event types: skipping exons (SE), alternative 5′/3′splice sites (SS), mutually exclusive exons (MX), retained introns (RI) and alternative first/last exons (FL) (https://github.com/comprna/SUPPA) and followed by PSI quantification per event using the .ioe files generated and the formatted file containing `salmon`-based TPMs. To study differential AS on `SUPPA2`'s generated events, `diffSplice` was applied with .psi and isoform TPM files split between ESC and Neurons with parameters `-m empirical` (to calculate significance), `--gene-correction` (to apply multiple testing correction by gene) and `-save-tpm-events`. Differential splicing results provided by `SUPPA2`, comprising 79 983 events, were filtered in R to keep each event for which the cognate transcript had TPM quantification available and the dPSI was not a missing value.

`rMATS` version turbo 4.1.0 was run on FASTQ files split between ESC and Neurons groups, using GENCODE M25 mouse genome annotation to build the `STAR` binary index and as GTF file. Differential splicing results were filtered in R to consider only exon skipping events without PSIs with missing values.

`rMATS` intermediate sorted BAM files generated were indexed using `samtools index` such that they could be used with the GENCODE M25 transcriptome annotation GFF3 file to build local splicing variation graphs using `MAJIQ builder`, followed by `MAJIQ psi` run to calculate PSI values and `MAJIQ deltapsi` to compare those PSIs between ESC and Neuron samples.

`Whippet` was used to build an index for AS events on which PSIs were quantified and compared between ESC and Neurons groups. `whippet-index` was run using FASTA genome and GTF gene annotation files from the GENCODE M25 annotation with the --suppress-low-tsl option to consider only well-supported transcript annotations (TSL2+) when building the splice graph. Inclusion levels, their confidence, and total reads for each defined node were quantified with `whippet-quant` on the FASTQ file pair for each sample under study. `whippet-delta` was then run to compare the generated psi.gz files, one for each sample, grouped into ESCs and Neurons. The psi.gz files and the differential splicing output file were filtered in R to consider only exon skipping ("CE, or core exon") events without missing PSI values.

For the between-tools comparison of differential AS splicing of the same sequences between mouse ESC and cortical glutamatergic neurons, we focused on the exon skipping events that behave as "alternative" (i.e., 0 < PSI < 1) in all ESC and Neuron samples. Such filtering criteria returned 738 events in `vast-tools`, 3 388 in `SUPPA2` and 1 909 in `rMATS` (Supplementary Figure 7). `MAJIQ` was not considered for the comparison between tools as the PSI quantifications for their units of local splicing variation (LSV) were not directly comparable with the exon skipping events considered for the other tools. Although `Whippet` also considers an annotation structure, in which PSIs are associated with the nodes of contiguous splice graphs, that is not directly comparable with those of other tools, we were able to compare `whippet-delta`'s differential splicing results for a subset of 367 exon skipping events associated with "core exon" nodes whose annotated alternative sequence coordinates' match VAST-DB's (Supplementary Figure 7).

To match inclusion quantifications of the same event across VAST-DB's, `SUPPA2`'s, `rMATS`', and `Whippet`'s annotations, we used `vast-tools`' coordinates of the annotated events in the format `chrN:exonStart-exonEnd` (available in column "COORD" of the output table) and adapted the function `parseSuppaAnnotation()` from `psichomics` (13). All processing was performed in the R environment for statistical computing.

#### `betAS` applied to comparisons between multiple sequential groups

Inspired by ANOVA, we extended the `betAS` pipeline to test differential AS between more than two groups. The coverage-dependent dispersion of values emitted from a beta distribution is a surrogate for the PSI uncertainty for each sample. Such dispersion can be compared within samples from the same group and compared with that found between groups. Thus, for each event, we consider within as the set of differences between pairs of samples that are part of the same group and between as the set of differences between pairs of samples from different groups.

Comparisons between the median absolute values of between and within are used to obtain an "F-like" statistic, which compares inter- and intra-group variabilities:

$$F_{stat} = \frac{\text{variation } \mathbf{between} \text{ samples}}{\text{variation } \mathbf{within} \text{ samples}} = \frac{\text{median}(|\mathbf{between}|)}{\text{median}(|\mathbf{within}|)}$$

For experimental designs with groups following a sequential relationship (e.g., time course), between distances are calculated by `betAS` in a sequential mode. Thus, sequential between are the set of between-group distances for any pair of consecutive groups in the sequence. Considering directional differences between groups (i.e., from each group to its predecessor), the modes of this set of differences will be associated with the prevalence of increasing (if positive) or decreasing (when negative) steps of a given magnitude, from which the monotonicity of change in inclusion levels can be empirically estimated. To accommodate this, we define, for each event, its monotonicity coefficient:

$$\text{monotonicity} = \frac{\#\,(\textbf{seq. differences}\text{ that are} > 0) - \#\,(\textbf{seq. differences}\text{ that are} < 0)}{\#\text{ all sequential }\textbf{differences}}$$

$$\text{monotonicity} = \frac{\#\,(\textbf{steps}\text{ that are} > 0) - \#\,(\textbf{steps}\text{ that are} < 0)}{\#\text{ all }\textbf{steps}}$$

$$\text{with }\textbf{steps} = \text{emitted values}_{\text{group } i} - \text{emitted values}_{\text{group } i\text{-}1}$$

Moreover, we extended the application of the $P_{diff}$ described in previous sections to calculate the probability that |between| > |within|:

$$P_{x=0} = \frac{\#\,(\text{differences }|\textbf{between}| - |\textbf{within}|\text{ that are} > 0)}{\#\text{ minimum length of between and within differences}}$$

$$P_{diff} = |P_{x=0} - 0.5| + 0.5$$
